# WDR60-mediated dynein-2 loading into cilia powers retrograde IFT and transition zone crossing

**DOI:** 10.1101/2021.09.19.459328

**Authors:** Ana R. G. De-Castro, Diogo R. M. Rodrigues, Maria J. G. De-Castro, Neide Vieira, Cármen Vieira, Ana X. Carvalho, Reto Gassmann, Carla M. Abreu, Tiago J. Dantas

## Abstract

The dynein-2 motor complex drives retrograde intraflagellar transport (IFT), playing a pivotal role in the assembly and functions of cilia. However, the mechanisms that regulate dynein-2 motility remain poorly understood. Here, we identify the *Caenorhabditis elegans* WDR60 homolog (WDR-60) and dissect the roles of this intermediate chain using genome editing and live imaging of endogenous dynein-2/IFT components. We find that loss of WDR-60 impairs dynein-2 recruitment to cilia and its incorporation onto anterograde IFT trains, reducing the availability of the retrograde motor at the ciliary tip. Consistently, we show that less dynein-2 motors power WDR-60-deficient retrograde IFT trains, which move at reduced velocities and fail to exit cilia, accumulating on the distal side of the transition zone. Remarkably, disrupting the transition zone’s NPHP module almost fully restores ciliary exit of underpowered retrograde trains in *wdr-60* mutants. This work establishes WDR-60 as a major contributor to IFT and the NPHP module as a roadblock to dynein-2 passage through the transition zone.

## INTRODUCTION

Cilia are microtubule-based structures that project outwards from the surface of most mammalian cell types. These antenna-like structures can produce mechanical force for locomotion or fluid flow (such as in the multiciliated airway epithelia), or sense extracellular signals that modulate developmental pathways, ultimately regulating cell proliferation and differentiation (Drummond, 2012). Regardless of their type, the assembly and functions of cilia depend on a bidirectional transport system known as intraflagellar transport (IFT) (Kozminski et al., 1993; Prevo et al., 2017; Webb et al., 2020). IFT is driven by two classes of molecular motors that travel on the ciliary microtubules that compose the axoneme. Kinesin-2 motors cooperate with both IFT-B and IFT-A complexes to power the transport of cargos in the anterograde direction from the base to the tip of cilia (Kozminski et al., 1995; Prevo et al., 2017). On the opposite direction, cytoplasmic dynein-2 (hereafter referred to as dynein-2) motors associate with the IFT-A complex to drive retrograde transport (Pazour et al., 1999; Porter et al., 1999; Wicks et al., 2000), which is critical for the retrieval of signaling molecules and the recycling of ciliary proteins (Prevo et al., 2017; Webb et al., 2020).

Between the base and the axoneme of cilia, a specialized ciliary gate, known as the transition zone (TZ), controls which proteins and membrane components enter and exit the cilium, thus isolating the ciliary environment from the cytoplasm (Garcia-Gonzalo and Reiter, 2017). The assembly and gating function of the TZ is a complex process that involves many soluble and membrane-bound components organized into the MKS (Meckel-Gruber syndrome) and NPHP (Nephronophthisis) modules (Garcia-Gonzalo and Reiter, 2017). These two modules cooperate for the building of Y-shaped structures (Y-links) that connect the proximal end of the axonemal doublets to the ciliary membrane at the region of the ciliary necklace (Blacque and Sanders, 2014). At the center of the TZ lies an apical ring (or central cylinder) that also contributes to the integrity and gating of the TZ (Li et al., 2016; Schouteden et al., 2015). Interestingly, dynein-2 has been recently shown to be required for the stability of the TZ (Jensen et al., 2018; Vuolo et al., 2018).

Problems in IFT or in the integrity of the TZ result in defects in cilia assembly or cilia-mediated functions that can lead to congenital developmental diseases, collectively known as ciliopathies (Drummond, 2012; Garcia-Gonzalo and Reiter, 2017). Mutations in genes coding for dynein-2 subunits are associated with severe skeletal dysplasias such as Jeune Syndrome, Asphyxiating Thoracic Dystrophy, Ellis-van Creveld Syndrome and Short-Rib Polydactyly Syndrome (SRPS), which in many cases is incompatible with fetal survival (Cossu et al., 2016; Dagoneau et al., 2009; Huber et al., 2013; McInerney-Leo et al., 2013; Merrill et al., 2009; Niceta et al., 2018; Schmidts et al., 2015; Schmidts et al., 2013; Taylor et al., 2015). How mutations in dynein-2 subunits lead to these disorders remains poorly understood.

Dynein-2 is a giant (>1 MDa) motor protein complex composed of heavy (HC), intermediate (IC), light intermediate (LIC) and light chains (LCs) (Toropova et al., 2019). A homodimer of two heavy chains (DHC2 encoded by *DYNC2H1*) comprises the core of the motor complex. Each DHC2 has an N-terminal (NT) tail that serves as a platform for the binding of other subunits, and a C-terminus (CT) comprising six AAA ATPase domains and a microtubule-binding domain that enable dynein-2 movement on microtubules. Two copies of the dynein-2-specific LIC3 (encoded by *DYNC2LI1*) bind and serve to stabilize the DHC2s (Mikami et al., 2002; Taylor et al., 2015; Toropova et al., 2019). The DHC2-LIC3 subcomplex binds an heterodimer of ICs, composed of WDR60 and WDR34 (encoded by *DYNC2I1* and *DYNC2I2*, respectively), through their CT β-propeller domains (Asante et al., 2014; Patel-King et al., 2013; Rompolas et al., 2007; Toropova et al., 2019). In turn, the WDR60-WDR34 heterodimer is stabilized through the binding of multiple LCs at their NT (Hamada et al., 2018; Schmidts et al., 2015; Toropova et al., 2019; Tsurumi et al., 2019).

Recent work analyzing the effects of depleting or disrupting the WDR60/WDR34 ICs in human cells has yielded inconsistent results, particularly regarding the requirement of WDR60 for ciliogenesis and cilia axoneme length control (Asante et al., 2014; Hamada et al., 2018; McInerney-Leo et al., 2013; Vuolo et al., 2018). Importantly, the impact of WDR60 loss on dynein-2 activity and dynamics during IFT has not been determined. This in part is due to the difficulty in visualizing and quantifying IFT kinetics of dynein-2 subunits (especially of DHC2) inside cilia of cultured cells (Hamada et al., 2018; Taylor et al., 2015; Vuolo et al., 2018).

In *Caenorhabditis elegans*, GFP/RFP-tagged dynein-2 HC and LIC subunits (see **Table S1** for nomenclature) are readily detectable inside cilia and easy to track during IFT (Mijalkovic et al., 2017; Schafer et al., 2003; Yi et al., 2017). Another advantage of this model is that mutations in dynein-2 subunits that are lethal in mice (Huangfu and Anderson, 2005; May et al., 2005; Rana et al., 2004; Wu et al., 2017) do not compromise viability in *C. elegans* (Schafer et al., 2003; Wicks et al., 2000; Yi et al., 2017). This is due to the fact that cilia are only present in *C. elegans* sensory neurons, which are analogous to human sensory cilia, such as those present in olfactory neurons. Despite being dispensable for viability, *C. elegans* cilia have key sensory functions that modulate easily quantifiable animal behaviors in response to environmental cues (Bae and Barr, 2008). These features make *C. elegans* a powerful model to dissect the roles of dynein-2 subunits during IFT.

Although most dynein-2 core subunits have been identified in *C. elegans*, clear homologs of WDR34 and WDR60 have remained unknown (Vuolo et al., 2020). Here, we identify the *C. elegans* WDR60 homolog (WDR-60) and dissect its contribution to ciliary recruitment of dynein-2 subunits, retrograde IFT, and cilia-mediated behavior. Using CRISPR/Cas9-mediated genome editing, we tagged endogenous WDR-60 with GFP and tracked its dynamics during IFT in cilia of sensory neurons. In addition, we generated a strain expressing a SRPS patient-equivalent WDR-60 truncation (McInerney-Leo et al., 2013) that lacks the DHC2-binding β-propeller domain (Toropova et al., 2019), and compared this mutant with a *wdr-60* null mutant. We show that WDR-60 is mostly dispensable for axoneme extension, but is required for efficient loading of dynein-2 onto anterograde IFT trains, for reaching maximum retrograde IFT velocity, and for dynein-2 crossing of the transition zone to exit cilia. By targeting specific TZ components, we were able to facilitate dynein-2 exit from WDR-60-deficient cilia, showing that dynein-2 is unable to overcome the resistance offered by the TZ barrier in the absence of WDR-60.

## RESULTS

### WDR-60 is recruited to cilia in *C. elegans* sensory neurons and undergoes IFT with similar kinetics as dynein-2 HC

We set out to identify the gene encoding the so far uncharacterized homolog of WDR60 in *C. elegans* (Hou and Witman, 2015; Vuolo et al., 2020). Through protein sequence alignments, we found that the *C27F2.1* gene in *C. elegans* encodes the protein with the highest sequence homology to human WDR60. Interestingly, *C27F2.1* (hereafter referred to as *wdr-60*) was one of the early candidate genes identified in a screen for transcripts specific for ciliated sensory neurons (Blacque et al., 2005). Like genes encoding for other dynein-2 subunits (Swoboda et al., 2000), *wdr-60* contains a predicted X-box sequence (**Figure S1A**), which is a target of the RFX-like transcription factor DAF-19 (Blacque et al., 2005).

To directly visualize and analyze the dynamics of the protein encoded by *wdr-60*, we used genome editing to introduce the coding sequence for a 3xFLAG::GFP tag at the endogenous *wdr-60* locus (**Figure S1A**). Similar to what has been described for dynein-2 LIC and HC (Schafer et al., 2003; Wicks et al., 2000), we found that WDR-60 expression is restricted to ciliated sensory neurons. To better define the tissue-specific expression of WDR-60, we performed the classic dye filling assay that takes advantage of a lipophilic fluorescent dye (DiI) that is specifically incorporated into ciliated sensory neurons that have their cilia in contact with the environment. As a control, we used a GFP knock-in strain of dynein-2 HC, GFP::CHE-3 (Yi et al., 2017). We found that the expression pattern of WDR-60::3xFLAG::GFP is identical to that of GFP::CHE-3, and perfectly matches the neurons that uptake dye (**Figure 1A**). While a large part of the signal is detected in the soma and dendrites of these neurons, WDR-60 is also found inside cilia, similar to what has been observed for GFP::CHE-3 (Yi et al., 2017).

**Figure 1.**
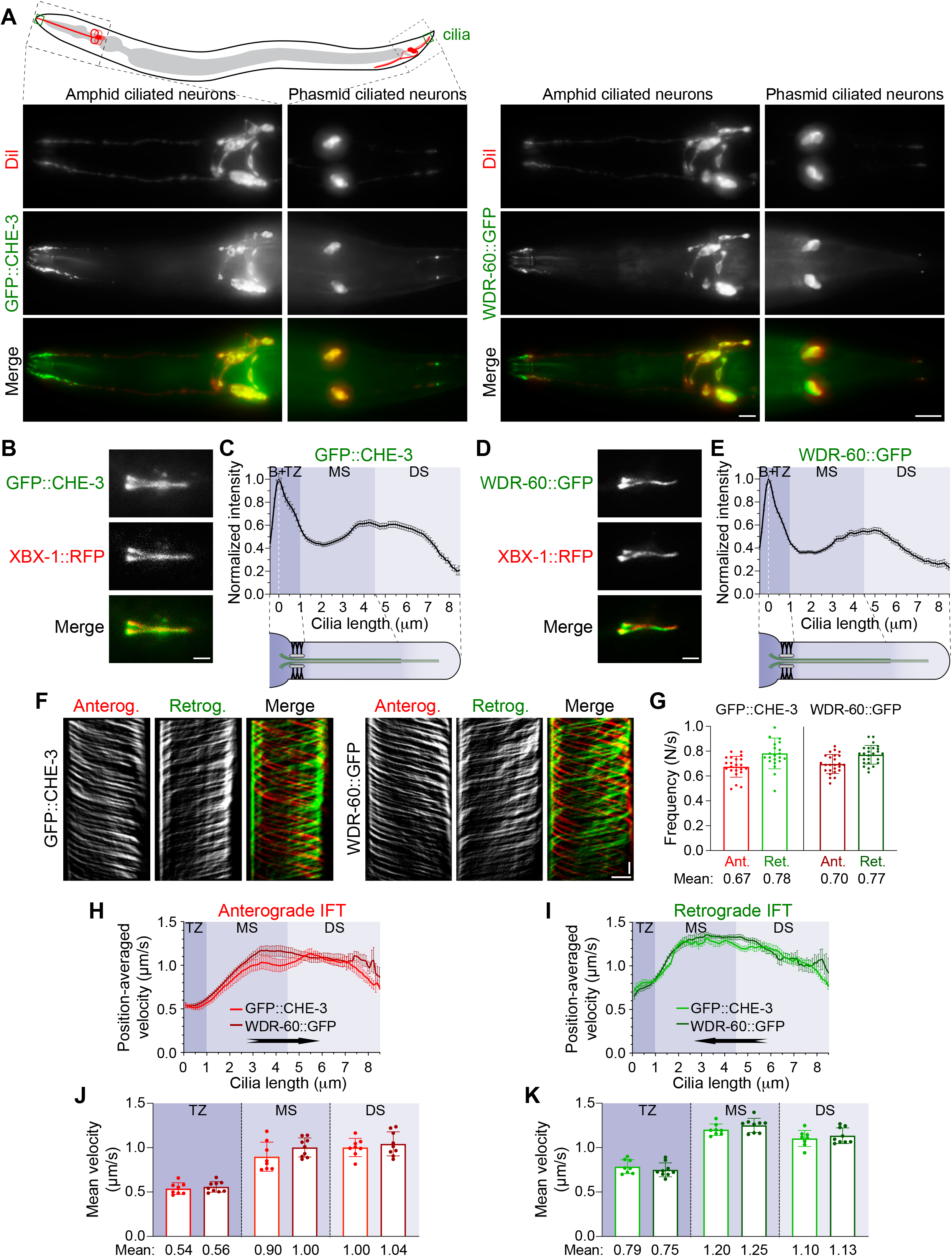
WDR-60 expression is restricted to ciliated sensory neurons, where it has similar distribution and IFT kinetics to the dynein-2 HC. **(A)** Endogenously-tagged WDR-60::3xFLAG::GFP is expressed in the same ciliated neurons that express dynein-2 HC (GFP::CHE-3). These are the same neurons that incorporate the DiI lipophilic dye. **(B)** Phasmid cilia co-expressing GFP::CHE-3 and XBX-1::RFP. **(C)** Quantification of GFP::CHE-3 signal intensity along cilia (N=109 cilia). **(D)** Phasmid cilia co-expressing WDR-60::3xFLAG::GFP and XBX-1::RFP. **(E)** Quantification of WDR-60::3xFLAG::GFP signal intensity along cilia (N=112 cilia). **(F)** Cilium kymographs of GFP::CHE-3 or WDR-60::3xFLAG::GFP. Single and merge channels for particles moving anterogradely and retrogradely are shown. **(G)** Mean IFT frequency of anterogradely and retrogradely moving GFP::CHE-3 and WDR-60::3xFLAG::GFP particles per second (N≥22 cilia). **(H,I)** Anterograde and retrograde velocities of GFP::CHE-3 and WDR-60::3xFLAG::GFP particles along cilia. **(J,K)** Mean velocities for each cilium sub-compartment (N≥430 particle traces were analyzed in ≥8 cilia). *B*, cilium base; *TZ*, transition zone; *MS*, middle segment; *DS*, distal segment. Scale bars: (A) 10 μm, (B,D) 2 μm, (F) vertical 5 sec, horizontal 2 μm.

When analyzing WDR-60::GFP ciliary distribution in more detail, we found that WDR-60 is particularly enriched at the ciliary base, as is the case for GFP::CHE-3 **(Figure 1B-E)**. Furthermore, both subunits co-localize with dynein-2 LIC, XBX-1::RFP (Yi et al., 2017). Next, we performed time-lapse imaging to gain insight into WDR-60 dynamics (**Movie S1**). We found that both anterograde and retrograde frequencies (**Figure 1F,G)** and velocities (**Figure 1H-K**) of WDR-60::GFP particles match those of GFP::CHE-3. Furthermore, retrograde WDR-60::GFP motility follows a triphasic model, as previously reported for GFP::CHE-3 (Yi et al., 2017) **(Figure 1I,K)**. Together, these data strongly support that *C27F2.1/wdr-60* encodes for the *C. elegans* WDR60 homolog, which undergoes IFT with kinetics that resemble those of dynein-2 HC.

### The β-propeller domain is important but not essential for WDR-60 incorporation into cilia

To determine the importance of WDR-60 for dynein-2-mediated IFT and cilia assembly, we first characterized WDR-60 levels and distribution in two distinct *wdr-60* mutants. We took advantage of the available *wdr-60* deletion allele *tm6453* (**Figure S1B**), a null mutation, and we engineered a *wdr-60* allele that produces a truncated form of WDR-60 specifically lacking the CT β-propeller domain (*ΔCT*; **Figure S1C; Figure 2A**), required for dynein-2 HC binding (Toropova et al., 2019). The *wdr-60(ΔCT)* mutant mimics a truncating mutation found in a SRPS patient (*WDR60*: c.1891C>T; p.Q631* (McInerney-Leo et al., 2013)).

**Figure 2.**
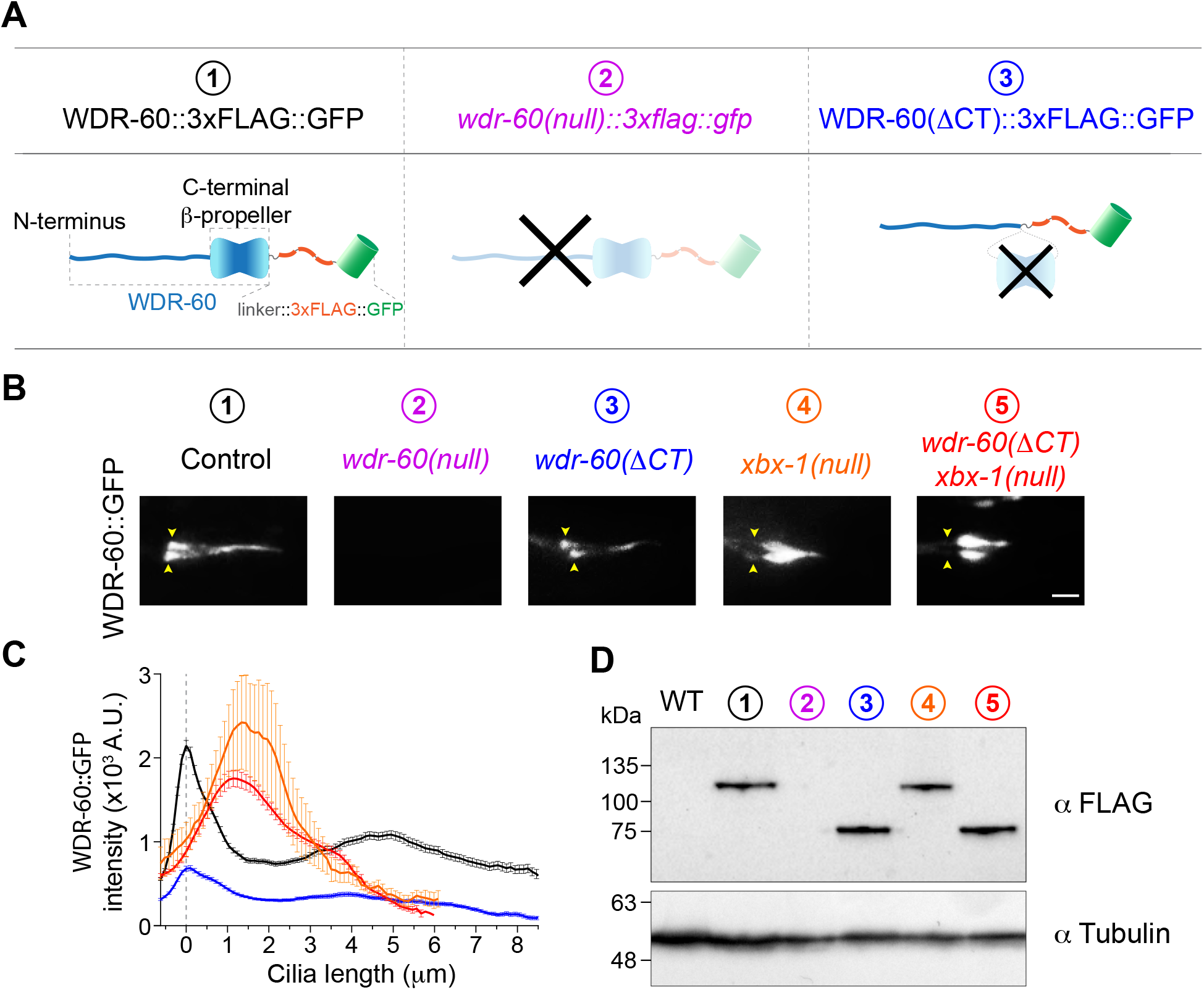
Truncation of the β-propeller domain reduces, but does not abolish, entry of WDR-60 into cilia. **(A)** Schematic representation of WDR-60 tagging with::3xFLAG::GFP in control, *wdr-60(tm6453)* null, and truncated *wdr-60* knock-in strains: (1) full length WDR-60, composed of an NT disordered region and a CT β-propeller domain (2) *wdr-60(tm6453)*, predicted to be a null mutant; (3) *wdr-60(ΔCT)*, expected to produce a protein composed of the WDR-60 NT fused to the::3xFLAG::GFP tag (lacking the β-propeller). **(B)** Phasmid cilia for each WDR-60 knock-in strain as indicated. Yellow arrowheads indicate the ciliary base. Note that no GFP signal is detected in the *wdr-60(tm6453)::3xflag::gfp* strain. Scale bar, 2 μm. **(C)** Quantification of GFP signal intensity distribution along the cilium in *wdr-60* mutants shown in B (N≥55 cilia). **(D)** Western blot of extracts from wild-type and WDR-60 knock-in strains using an anti-FLAG antibody. The predicted sizes are 105.6 kDa for WDR-60::3xFLAG::GFP and 63.2 kDa for WDR-60(ΔCT)::3xFLAG::GFP truncation. No signal is detected in *wdr-60(tm6453)::3xflag::gfp* extracts, demonstrating that this is indeed a null strain. α-Tubulin was used as a loading control.

As endogenous labelling of WDR-60 did not alter IFT kinetics (**Figure 1G-K**), we inserted the same 3xFLAG::GFP tag sequence in frame with the 3’ end of both *wdr-60* mutants (**Figure S1A, Figure 2A**). Together with the dye-filling assay, this allowed us to analyze the overall integrity of cilia while comparing the neuronal localization and relative levels of WDR-60 in wildtype and in each mutant (**Figure S2**). Interestingly, and in contrast to the null *xbx-1(ok279)* mutant, both *wdr-60* mutants had all ciliated sensory neurons stained with DiI, suggesting that sensory cilia can form and uptake dye. No GFP signal was detectable in sensory neurons or in cilia of the *wdr-60(tm6453)* mutant, indicating that no WDR-60 is produced in this strain. In contrast, the GFP signal in neurons of the *wdr-60(ΔCT)* mutant was readily visible and overlapped with the neuronal pattern of the dye (**Figure S2; Figure 2A,B**). Interestingly, the ciliary signal of WDR-60(ΔCT)::GFP was overall weaker than in controls (~3-fold reduction; **Figure 2C**) but showed a similar distribution profile along the axoneme, suggesting that a fraction of WDR-60(ΔCT) is able to enter cilia and undergo IFT. Strikingly, the absence of dynein-2 LIC, which is known to destabilize dynein-2 HC (Taylor et al., 2015; Yi et al., 2017), did not significantly affect the ciliary recruitment of WDR-60::GFP or WDR-60(ΔCT)::GFP (**Figure 2B,C)**. However, it did lead to the accumulation of both forms of WDR-60 inside cilia, likely due to the complete block of retrograde IFT that occurs in the *xbx-1(null)* mutant (Schafer et al., 2003; Yi et al., 2017). We conclude that WDR-60 can be recruited to cilia independently of dynein-2 LIC and HC subunits.

Taking advantage of the 3xFLAG epitope in our tag, we performed immunoblotting to determine whether the reduction in ciliary levels of mutant WDR-60 reflected differences in overall protein levels (**Figure 2D**). Given that no protein bands were detectable in *wdr-60(tm6453)* worm extracts, we conclude that this mutant strain is indeed a *wdr-60* null. In contrast, the levels of WDR-60(ΔCT)::GFP were comparable to those of full-length WDR-60::GFP, indicating that the reduced ciliary recruitment of WDR-60(ΔCT)::GFP is due to loss of the CT β-propeller rather than a downregulation of protein levels.

### Disruption of WDR-60 reduces dynein-2 loading into cilia and the kinetics of retrograde IFT

Loss of the dynein-2 LIC XBX-1 destabilizes the dynein-2 HC CHE-3, completely abolishing its recruitment to cilia, blocking retrograde IFT and, consequently, axoneme extension. The resulting cilia are severely truncated and bulged (Schafer et al., 2003; Yi et al., 2017). To directly assess the impact of WDR-60 disruption on cilia and other dynein-2 subunits, we crossed the *wdr-60* mutants with knock-in strains of GFP::CHE-3/XBX-1::RFP (Yi et al., 2017) and analyzed their ciliary recruitment and distribution. While both *wdr-60* mutants were capable of assembling seemingly normal cilia (with only a minor reduction in length in the *wdr-60* null mutant), we observed a strong reduction in the total levels of ciliary GFP::CHE-3 (~40%; **Figure 3A-C**). Interestingly, we also found that the remaining pool of WDR-60-deficient dynein-2 accumulated particularly near the ciliary base (**Figure 3D**). Considering that GFP::CHE-3 levels were not greatly altered in the soma of the ciliated phasmid neurons of *wdr-60* mutants (**Figure S2C**), these observations uncover that WDR-60 contributes to both recruitment and ciliary distribution of dynein-2.

**Figure 3.**
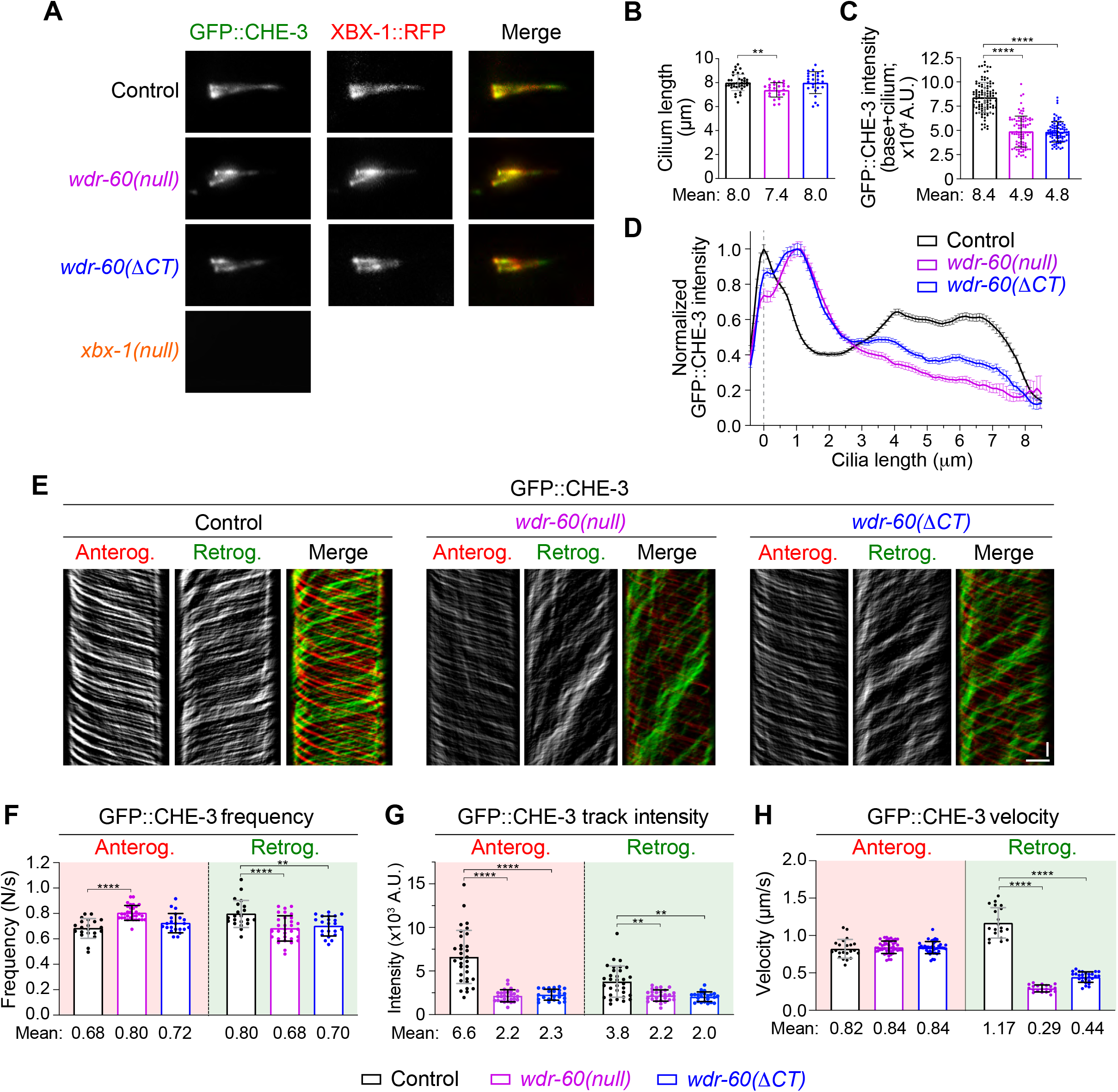
*wdr-60* mutants have reduced dynein-2 recruitment and incorporation into cilia, accompanied by impaired retrograde IFT. **(A)** Phasmid cilia co-expressing GFP::CHE-3 and XBX-1::RFP. **(B)** Cilia length in *wdr-60* mutants (N≥24 cilia). **(C)** Total signal intensity of GFP::CHE-3 from the base to the tip of cilia and **(D)** relative distribution of GFP::CHE-3 along cilia (N≥38 cilia). **(E)** GFP::CHE-3 kymographs of phasmid cilia of the indicated strains. Single and merge channels for particles moving anterogradely and retrogradely are shown. **(F)** Frequency of IFT particles detected at the distal segment of cilia (N≥20 cilia). **(G)** Quantification of the average intensity of GFP::CHE-3 particles moving on anterograde and retrograde tracks (N≥345 particle traces were analyzed in ≥23 cilia). **(H)** Velocity of anterograde and retrograde GFP::CHE-3 particles in control and *wdr-60* mutants (N≥300 particle traces were analyzed in ≥20 cilia). Scale bars: (A) 2 μm, (E) vertical 5 sec, horizontal 2 μm.

To determine when these WDR-60-associated phenotypes start manifesting and whether they vary with aging, we repeated our analysis of GFP::CHE-3 recruitment and distribution in developing and post-adulthood animals. We found that, at as early as the larval stage 2 (L2) of the *wdr-60(null)* mutant, the ciliary levels of GFP::CHE-3 were already reduced and its distribution altered, albeit less than in young adults. This suggests that WDR-60-associated dynein-2 phenotypes arise early on, but get progressively worse as the mutant animals develop (**Figure S3**). In addition, we found that the abnormal distribution of GFP::CHE-3 does not significantly change with age in *wdr60(null)* animals (7 and 18 days post-adulthood, **Figure S3C-H**), suggesting that there is no age-dependent suppression of these WDR-60-associated phenotypes, contrasting to what has been observed for some IFT mutants (Cornils et al., 2016).

To gain further insight into the importance of WDR-60 for dynein-2 loading and dynamics inside cilia, we analyzed the IFT kinetics of GFP::CHE-3 by time-lapse imaging (**Figure 3E-H, Movie S2**). While we found a small increase in the frequency of GFP::CHE-3 particles moving in the anterograde direction in the *wdr-60(null)* mutant, the number of particles in the retrograde direction was significantly reduced in both mutants (~15%; **Figure 3F**). Importantly, we found that both the loss of WDR-60 and the truncation of its β-propeller led to a strong reduction in the average amount of GFP::CHE-3 transported on anterograde tracks (~67%; **Figure 3G**). This establishes a role for WDR-60 in the loading of dynein-2 onto anterograde IFT trains. Consistent with this, we found that the intensity of GFP::CHE-3 moving on retrograde tracks was significantly reduced in both *wdr-60* mutants, suggesting that each retrograde IFT train is being powered by less dynein-2 motors. In addition, while the velocity of anterograde trains carrying GFP::CHE-3 remained similar to controls, we observed a strong reduction in the velocity of GFP::CHE-3 driven retrograde trains in both *wdr-60* mutants (3 to 4-fold; **Figure 3H**).

Taken together, these results show that loss of WDR-60 or truncation of its β-propeller reduces the loading of dynein-2 HC onto anterograde IFT trains, and consequently the pool of dynein-2 available at the tip of cilia to power retrograde IFT. In agreement, less dynein-2 motors were present in particles moving retrogradely, which may explain the reduced kinetics of retrograde IFT in *wdr-60* mutants, and the inability of dynein-2 to fully return to the ciliary base.

### WDR-60 is required for efficient recycling of IFT components and contributes to cilia-mediated behavior

To better understand the importance of WDR-60 in IFT, we analyzed fluorescently-labeled subunits of the IFT-A/B complexes (**Figure 4A-F**) and anterograde kinesins (**Figure S4B,C**). Measurements of cilia expressing CHE-11::mCherry (IFT140) or IFT-74::GFP confirmed the small but significant decrease in cilium length in the *wdr-60(null)* mutant (**Figure 4A,B,D,E**). We note, however, that this minor defect in axoneme extension is distinct from the severely shortened and bulged cilia phenotype caused by XBX-1 loss (**Figure 4A,D**).

**Figure 4.**
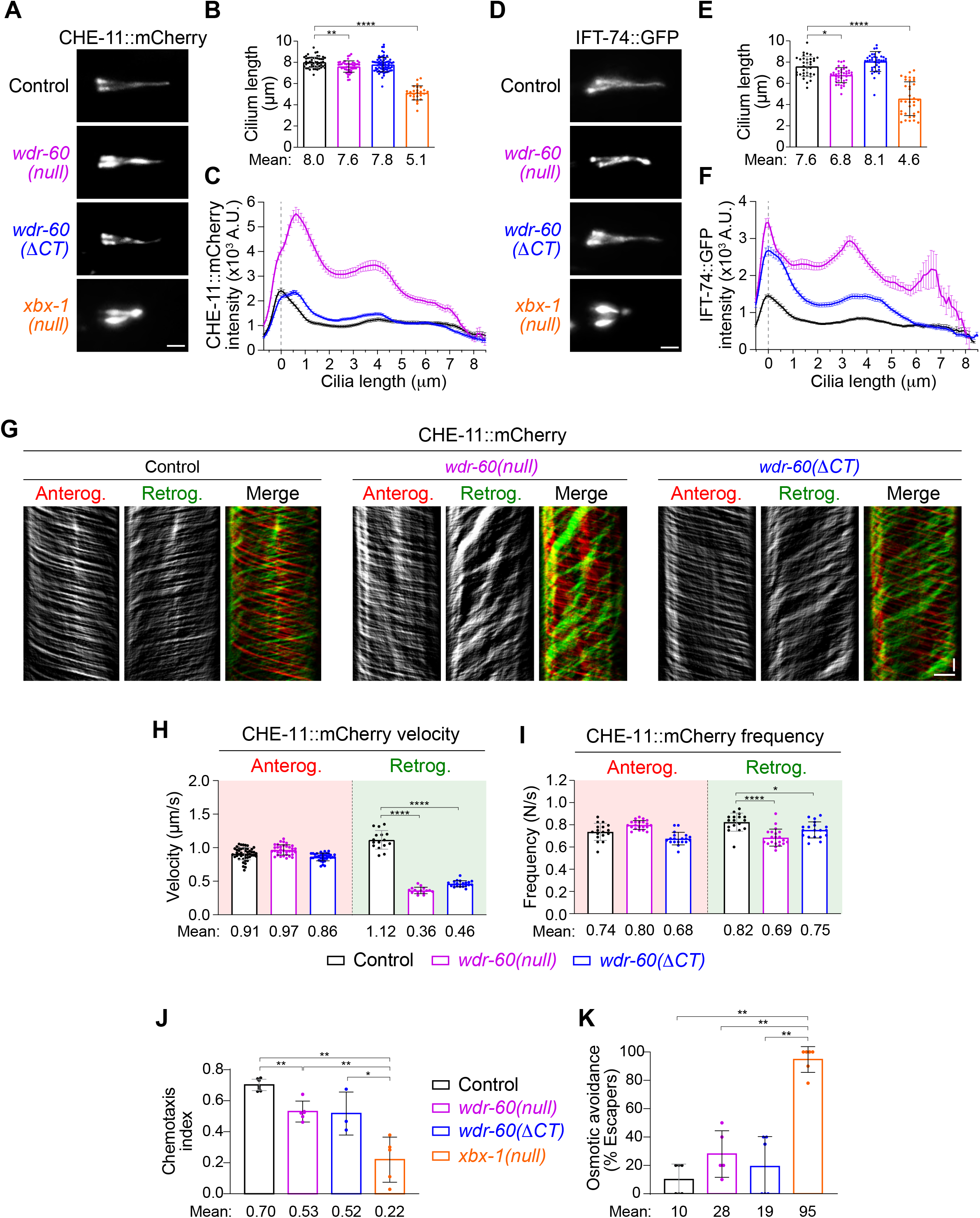
*wdr-60* mutants have reduced retrograde transport of IFT-A and IFT-B complexes, and lower efficiency of cilia-mediated signaling. **(A,D)** Phasmid cilia from control, *wdr-60(null), wdr-60(ΔCT)* and *xbx-1(null)* animals expressing (A) CHE-11::mCherry or (D) IFT-74::GFP. **(B,E)** Cilia length measured with (B) CHE-11::mCherry or (E) IFT-74::GFP (N≥26 and N≥34 cilia, respectively). **(C,F)** Quantification of the average intensity of (C) CHE-11::mCherry and (F) IFT-74::GFP along cilia (N≥42 and N≥66 cilia, respectively). **(G)** CHE-11::mCherry kymographs of phasmid cilia from the indicated strains. **(H)** Mean velocity of CHE-11::mCherry particles moving on anterograde and retrograde tracks (N≥225 particle traces were analyzed in ≥15 cilia). **(I)** Frequency of IFT particles detected at the distal segment of cilia (N≥17 cilia). **(J,K)** WDR-60 is required for efficient sensory cilia functions. **(J)** Chemotaxis index for the attractant isoamyl alcohol (N≥450 animals tracked over ≥3 assays). **(K)** Osmotic avoidance assay to test whether sensory cilia detect a hypertonic glycerol barrier (N≥20 animals tracked over ≥4 assays). The *xbx-1(null)* strain was used for comparison. Scale bars: (A,D) 2 μm, (G) vertical 5 sec, horizontal 2 μm.

When analyzing the ciliary distribution of CHE-11::mCherry, we found that this IFT-A component accumulated predominantly near the ciliary base in both *wdr-60* mutants (**Figure 4A,C**), similar to our observations with GFP::CHE-3. Interestingly, the total levels of CHE-11::mCherry retained inside cilia were substantially higher in *wdr-60(null)* mutant when compared with the *wdr-60(ΔCT)* mutant.

Consistent with defects in dynein-2 function, we found that the retrograde velocity of CHE-11 was strongly reduced in both *wdr-60* mutants (~3-fold; **Figure 4G,H, Movie S3**). In addition, the frequency of CHE-11::mCherry tracks was also significantly reduced in the retrograde direction (~16% lower in *wdr-60(null)* cilia; **Figure 4I**).

The IFT-B subunit IFT-74::GFP accumulated at multiple places along cilia in both *wdr-60* mutants (**Figure 4D,F, Figure S4A, Movie S4**), underscoring the importance of WDR-60 in dynein-2-mediated transport of the IFT-B machinery to the ciliary base. When analyzing the distribution of kinesins, we also observed ciliary accumulations for the kinesin-2-associated protein KAP-1 (KIFAP3) and, to a lesser extent, for the distal segment kinesin OSM-3 (KIF17) (**Figure S4B,C**). Altogether, these results show that loss of WDR-60 greatly impairs removal of IFT components from cilia.

To determine the impact of *wdr-60*-associated IFT defects on ciliary functions, we analyzed cilia-dependent behavior in our mutant strains. We tested chemotaxis attraction to isoamyl alcohol and osmotic tolerance to high concentrations of glycerol (**Figure 4J,K**). Both *wdr-60* mutants showed modest defects in these assays, contrasting with the *xbx-1(null)* mutant, in which chemotaxis attraction and osmotic tolerance is severely compromised. These results suggest that although WDR-60 plays critical roles in dynein-2-mediated IFT, WDR-60-deficient sensory cilia remain partially functional.

### TZ integrity and gating function are maintained in *wdr-60* mutants but not in the *xbx-1* mutant

Given that dynein-2 and IFT-A components were recently shown to be required for maintaining the TZ barrier (Jensen et al., 2018; Scheidel and Blacque, 2018; Vuolo et al., 2018), we investigated whether the integrity and gating capacity of the TZ are affected in *wdr-60* and *xbx-1* mutants. We analyzed the localization of four TZ components: TMEM-107, NPHP-4, MKS-6 and MKSR-1 (Jensen et al., 2018; Lambacher et al., 2016; Prevo et al., 2015; Schouteden et al., 2015; Williams et al., 2011). In agreement with what has been reported for CHE-3 and IFT-A mutants (Jensen et al., 2018; Scheidel and Blacque, 2018), we found that loss of XBX-1 results in ectopic localization of these TZ components along the ciliary axoneme (**Figure 5**). In contrast, the localization of TZ components in *wdr-60* mutants was indistinguishable from controls (**Figure 5**), suggesting that loss of WDR-60 does not affect the integrity of the TZ. To directly test the integrity and the gating capacity of the TZ in *wdr-60* mutants, we analyzed its ability to block the entry of RPI-2::GFP (RP2), a component of the periciliary membrane compartment, restricted to the base of cilia (Jensen et al., 2018). While loss of XBX-1 resulted in abnormal entry of RPI-2::GFP into cilia, no RPI-2::GFP signal was detectable inside cilia in *wdr-60* mutants (**Figure 5C,D**). These results strongly attest that the loss of WDR-60 does not compromise TZ integrity nor its gating function in *C. elegans*. In addition, our results uncover an important role for the dynein-2 LIC XBX-1 in maintaining the TZ barrier, likely by stabilizing CHE-3.

**Figure 5.**
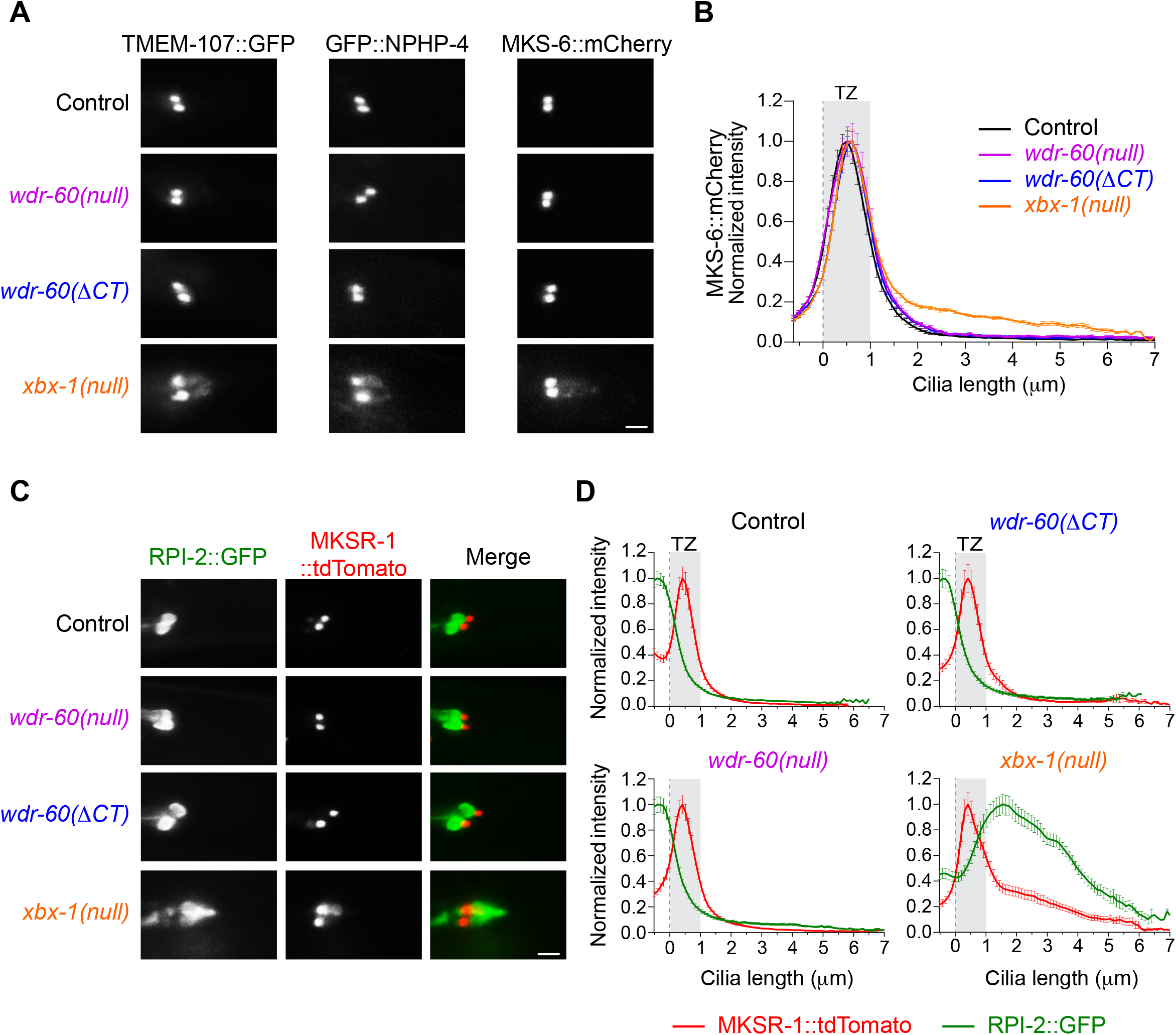
The integrity and gating function of the TZ are maintained in *wdr-60* mutants but compromised in the *xbx-1* mutant. **(A)** Analysis of the confinement of several components of the MKS (TMEM-107::GFP and MKS-6::mCherry) and the NPHP modules (GFP::NPHP-4) of the TZ in phasmid cilia of the indicated strains. **(B)** Quantification of MKS-6::mCherry signal intensity confined at the TZ and dispersed along cilia (N≥38 cilia). **(C)** Relative localization of the non-ciliary membrane protein RPI-2::GFP to the TZ (labeled with MKSR-1::tdTomato) in phasmid cilia of the indicated strains. **(D)** Signal overlap between these components and quantification of the amount of RPI-2::GFP leaking into cilia (N≥33 cilia). Gray rectangles highlight the TZ region, defined by MKS-6 or MKSR-1 localization. Scale bars, 2 μm.

### WDR-60 is required for dynein-2 passage through the TZ to exit cilia

To more precisely determine where the ciliary pool of dynein-2 accumulates in *wdr-60* mutants, we co-labeled GFP::CHE-3 with markers specific for the ciliary base (mCherry::HYLS-1 (Schouteden et al., 2015)) and for the TZ (MKS-6::mCherry (Williams et al., 2011)). In controls, GFP::CHE-3 mainly accumulates at the ciliary base (**Figure 6A**), before entering cilia through the TZ. In contrast, we found that GFP::CHE-3 accumulates mostly at the distal side of the TZ in both *wdr-60* mutants (**Figure 6B,C**).

**Figure 6.**
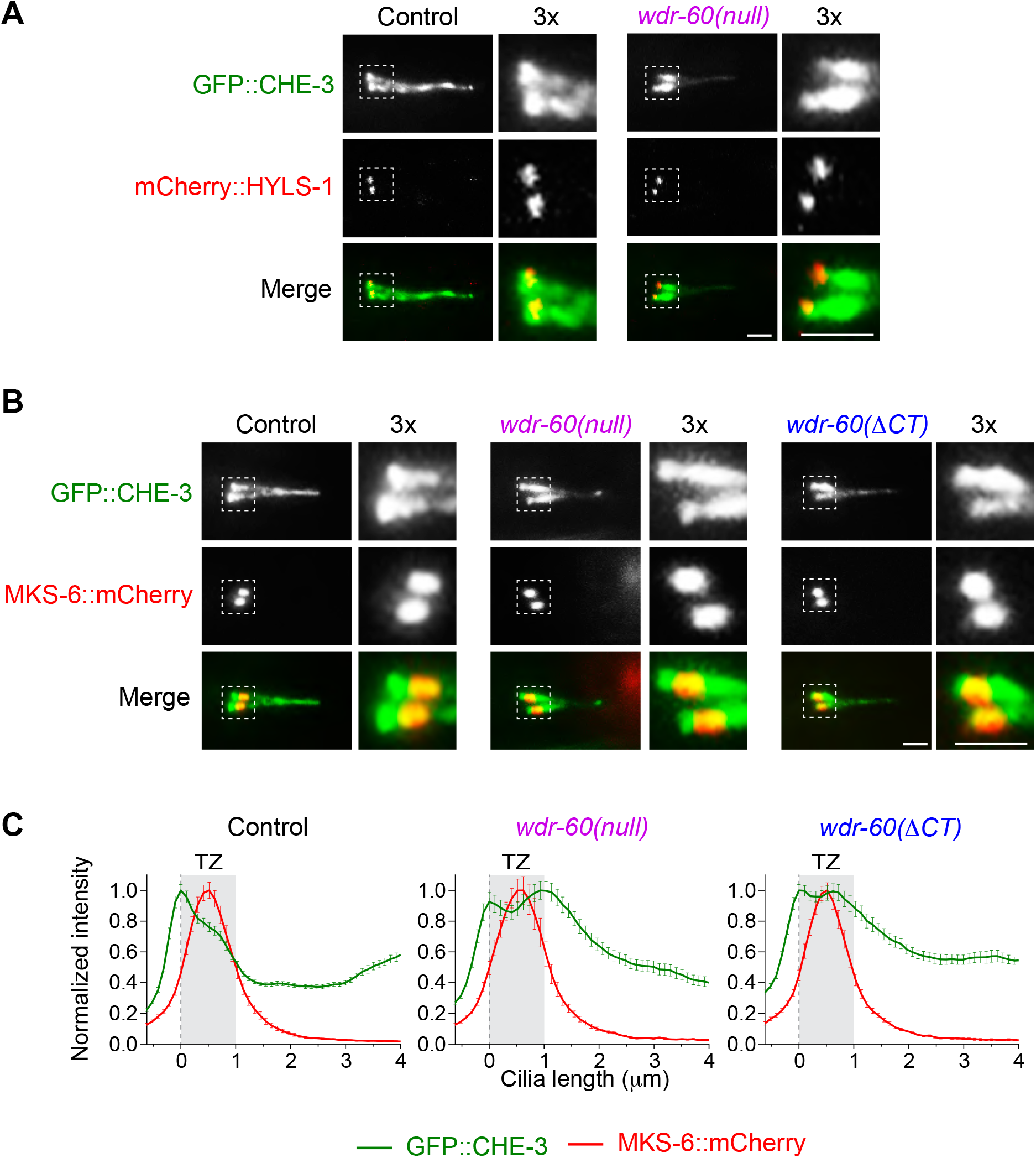
Dynein-2 accumulates on the distal side of the TZ, unable to complete retrograde IFT. **(A)** GFP::CHE-3 localization relative to the centriolar wall component mCherry::HYLS-1 at the base of phasmid cilia of the indicated strains. **(B)** GFP::CHE-3 localization relative to the MKS-6::mCherry TZ component in phasmid cilia of the indicated strains. **(C)** Quantification GFP::CHE-3 signal distribution in relation to the TZ, as determined by MKS-6 localization (N≥38 cilia). Gray rectangles define the TZ region. 3x magnifications of the square section in each micrograph were included to better visualize the distribution of the dynein-2 HC relative to the base and TZ of *wdr-60* mutant cilia. Scale bars, 2 μm.

Recent studies have shown that retrograde trains slow down as they cross the TZ, suggesting that this ciliary gate offers resistance to the passage of retrograde IFT trains (Jensen et al., 2015; Oswald et al., 2018; Prevo et al., 2015). Given that loss of WDR-60 reduces the amount of dynein-2 driving retrograde trains and impairs retrograde IFT velocity (**Figure 3)**, we hypothesized that WDR-60-deficient retrograde trains may not be able to generate enough force to push through the TZ barrier, and are consequently unable to exit cilia. As removal of MKS-5 (RPGRIP1L), a key component for the assembly of all TZ structures, significantly increases the velocity of IFT trains moving in the TZ region (Jensen et al., 2015), we reasoned that the exit of WDR-60-deficient dynein-2 from cilia might be facilitated by disrupting MKS-5. In agreement, we found that GFP::CHE-3 no longer accumulated on the distal side of the TZ in the *mks-5(tm3100);wdr-60(null)* double mutant (**Figure 7A-D**). Instead, the GFP::CHE-3 distribution profile in this mutant was similar to that observed in controls. This result further supports that retrograde trains driven by dynein-2 are unable to efficiently cross the TZ in *wdr-60* mutants.

**Figure 7.**
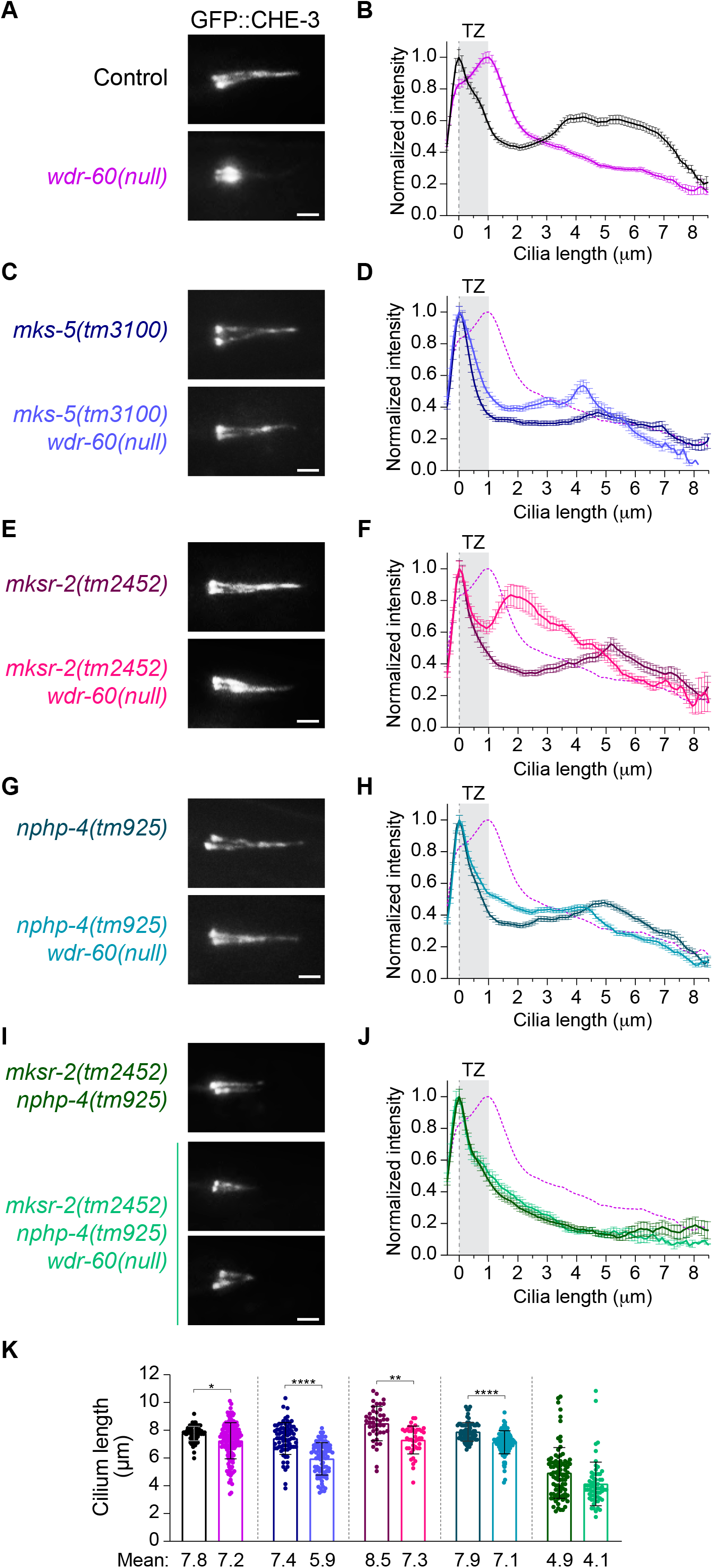
Disruption of specific TZ components rescues dynein-2 accumulation inside cilia of *wdr-60* mutants. **(A,C,E,G,I)** Representative examples of phasmid cilia of the indicated *wdr-60* and TZ mutant genotypes, expressing GFP::CHE-3. **(B,D,F,H,J)** Relative distribution of GFP::CHE-3 signal intensity along cilia. The purple dashed line represents the data from the *wdr-60(null)* in (B). Gray rectangles highlight the TZ, as previously defined. N≥108 cilia for (B), N≥74 cilia for (D), N≥42 cilia for (F), N≥80cilia for (H) and N≥64 cilia (J). **(K)** Length of the cilia analyzed in (B,D,F,H,J) from the same color-coded genotypes indicated in (A,C,E,G,I). Loss of WDR-60 combined with TZ mutations always resulted in a slight decrease of cilia length relative to the respective TZ mutant control. Scale bars, 2 μm.

To better dissect which TZ modules restrict dynein-2 passage, we next examined GFP::CHE-3 distribution in *wdr-60(null)* cilia after disrupting key components required for the assembly of each TZ module. Removal of the MKS module by inhibiting MKSR-2 (B9B2) or CEP-290 with the *mksr-2(tm2452)* or *cep-290(tm4927)* mutations did not prevent the accumulation of GFP::CHE-3 near the TZ region of *wdr-60(null)* cilia (**Figure 7E,F, Figure S5A,B**). In contrast, disrupting the NPHP module with the *nphp-4(tm925)* mutation in the *wdr-60(null)* background almost completely rescued GFP::CHE-3 accumulation at the distal side of the TZ (**Figure 7G,H**). Joint disruption of NPHP-4 and MKSR-2 further enhanced this rescue effect (**Figure 7I,J**), although we note that both cilia size and GFP::CHE-3 levels along cilia were strongly reduced in the *mksr-2(tm2452);nphp-4(tm925)* double mutant (**Figure 7K, Figure S5C,D**). We conclude that NPHP is the main module restricting the passage of underpowered retrograde trains through the TZ in *wdr-60(null)* cilia.

Next, we tested whether disrupting the TZ could compensate for more severe retrograde IFT defects, such as those caused by a mutation in the microtubule-binding domain of CHE-3 (K2935Q), that completely blocks dynein-2 motility and leads to severely truncated cilia (Yi et al., 2017). In contrast to the rescue that we observed earlier in the *wdr-60(null)* mutant background, disrupting MKS-5, NPHP-4 or both MKSR-2/NPHP-4 failed to prevent GFP::CHE-3(K2935Q) accumulation inside cilia (**Figure 8A-H**). These results show that even the complete removal of the TZ barrier is not sufficient to rescue the accumulation of non-motile dynein-2 inside cilia. Although cilia size slightly increased in the absence of MKS-5 or NPHP-4 (**Figure 8I**), none of the TZ mutants were able to restore anterograde IFT in animals expressing GFP::CHE-3(K2935Q) (**Figure 8J; Movie S5**). Thus, IFT requires a minimum amount of functional dynein-2 regardless of the state of the TZ barrier.

**Figure 8.**
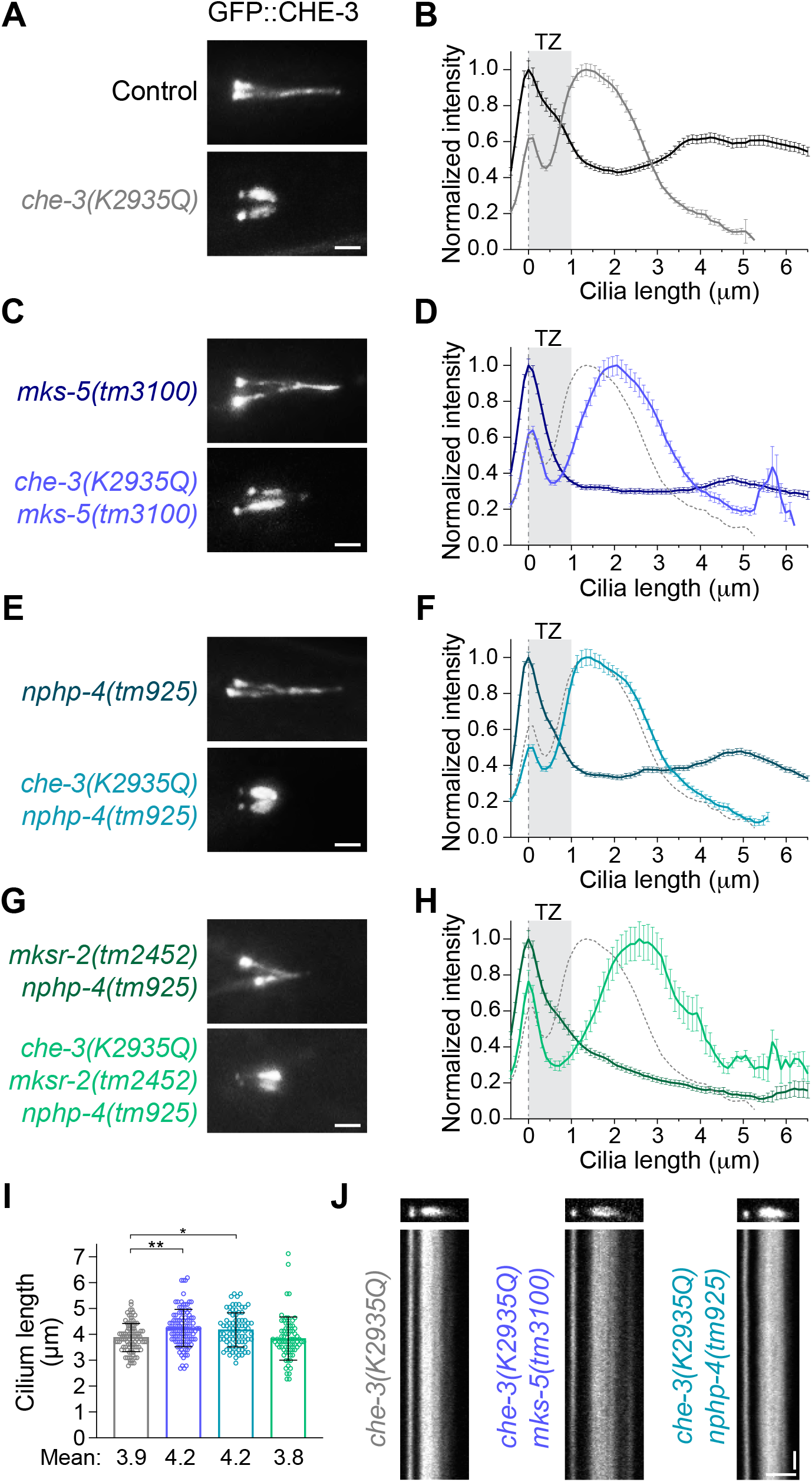
Complete loss of dynein-2 motility prevents rescue of ciliary accumulations even when the TZ barrier is completely disrupted. **(A,C,E,G)** Representative examples of phasmid cilia of the indicated TZ mutant genotypes, expressing wild-type GFP::CHE-3 or the non-motile GFP::CHE-3(K2935Q). **(B,D,F,H)** Relative distribution of GFP::CHE-3(K2935Q) signal intensity along cilia, in control and TZ mutant backgrounds. The gray dashed line represents the data from the GFP::CHE-3(K2935Q) mutant in (B). Gray rectangles highlight the TZ, as previously defined. N≥105 cilia for (B). N≥104 cilia for (D). N≥80 cilia for (F) N≥86 cilia (H). **(I)** Length of GFP::CHE-3(K2935Q) mutant cilia with the same color-coded genotypes indicated in (A,C,E,G) (N≥38 cilia). **(J)** Cilia and the respective kymographs from the specified strain genotypes. No anterograde or retrograde IFT was detectable in the GFP::CHE-3(K2935Q) mutant, not even in combination with the disruption of MKS-5 or NPHP-4. Scale bars: (A,C,E,G) 2 μm, (J) vertical 5 sec, horizontal 2 μm.

### The NPHP module restricts dynein-2 movement through the TZ

Next, we investigated whether the kinetics of WDR-60-deficient dynein-2 were altered by the complete removal of the TZ barrier or by the loss of the NPHP module. Consistent with a previous study (Jensen et al., 2015), we observed that loss of MKS-5 increases both the anterograde and retrograde velocities of GFP::CHE-3 particles in the TZ region (**Figure 9A,C,D, Movie S6**). We also observed a similar albeit more modest increase in IFT velocities in the TZ region in the *nphp-4* mutant background. Importantly, we found that loss of either MKS-5 or NPHP-4 increased the retrograde velocity of GFP::CHE-3 in the TZ region of WDR-60-deficient cilia (**Figure 9A,C,D, Movie S7**). Interestingly, retrograde IFT velocity in the middle segment of *wdr-60(null)* cilia also increased upon the removal of MKS-5 or NPHP-4 (**Figure 9C**), suggesting that clearing the accumulated IFT trains near the TZ allows for more steady buildup of retrograde IFT velocities in these mutants. We note that IFT frequency was not greatly affected by disruption of MKS-5 or NPHP-4 (**Figure 9B**), suggesting that loss of the TZ barrier does not compromise the poorly understood mechanisms regulating the rate of IFT injection into cilia.

**Figure 9.**
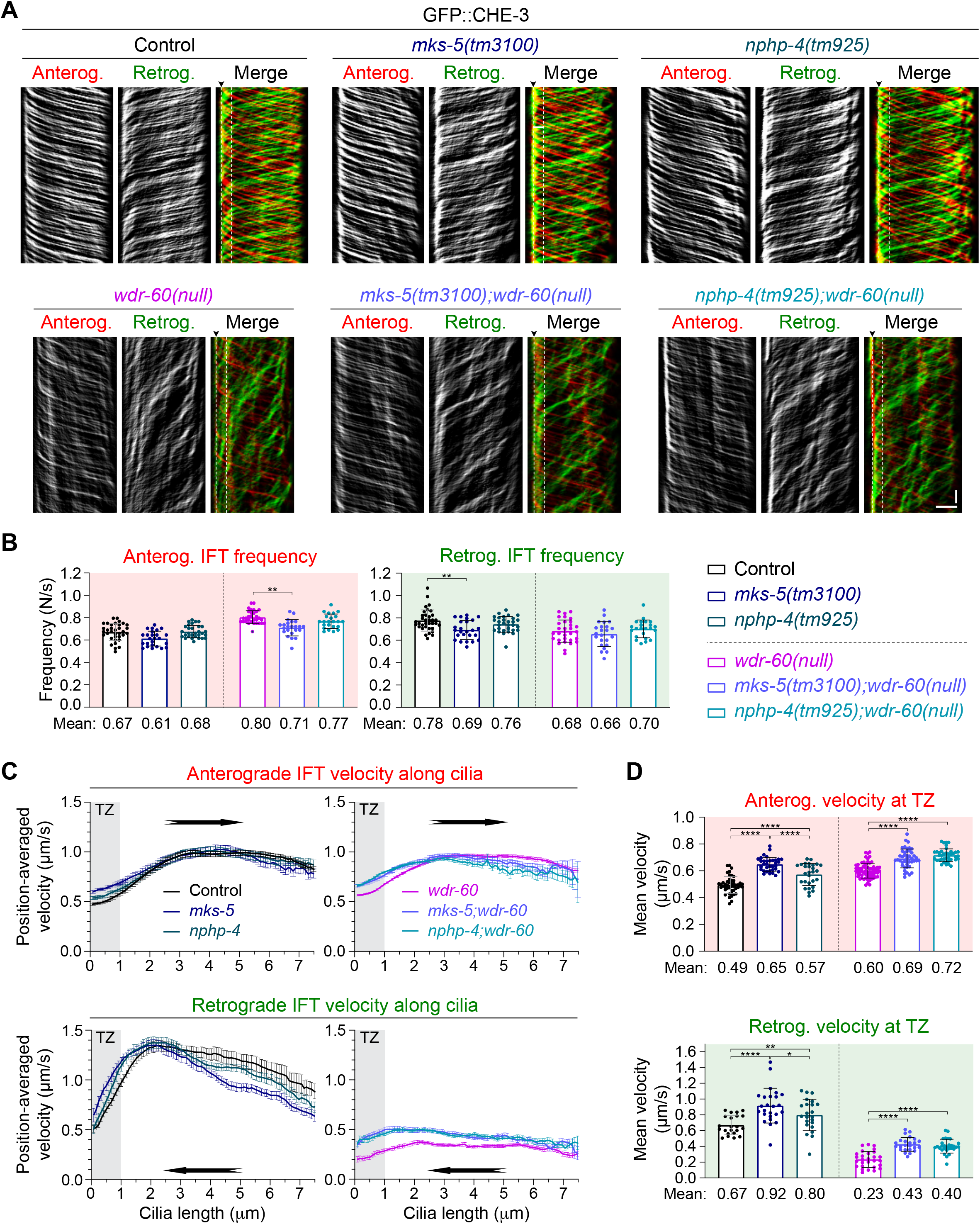
Disruption of NPHP-4 is sufficient to increase the velocity of WDR-60-deficient dynein-2 through the TZ region. **(A)** GFP::CHE-3 kymographs of phasmid cilia from the indicated strains. Single and merge channels for particles moving anterogradely and retrogradely are shown. Arrowhead labels the base, white dashed line marks the limit of the TZ region (1 μm). Scale bars, vertical 5 sec, horizontal 2 μm. **(B)** Frequency of IFT particles detected at the distal segment of cilia, for each represented strain (N≥23 cilia). **(C)** GFP::CHE-3 velocity moving in the anterograde and retrograde direction along the cilia of the represented genotypes (N≥345 particle traces were analyzed in ≥23 cilia for each strain). Gray rectangles highlight the TZ, as previously defined. **(D)** Average velocity of GFP::CHE-3 crossing the TZ region of the same cilia analyzed in (C), for each represented genotype.

Altogether, these results further support that WDR-60 loss impairs dynein-2 passage through the TZ, and that the NPHP module restricts dynein-2 exit from WDR-60-deficient cilia.

## DISCUSSION

### WDR-60 is incorporated into cilia even in the absence of dynein-2

Our data reveals that WDR-60 is specifically expressed in ciliated sensory neurons in *C. elegans* and undergoes IFT with kinetics similar to those reported for the dynein-2 HC (GFP::CHE-3)(Yi et al., 2017). Moreover, our findings indicate that WDR-60(ΔCT) is robustly expressed, showing that the β-propeller is not required for WDR-60 stability. Interestingly, we find that the WDR-60 NT on its own can be recruited to the cilium base and incorporated into IFT trains, albeit less efficiently than full-length WDR-60. Furthermore, dynein-2 HC destabilization through XBX-1 loss resulted in WDR-60 and WDR-60(ΔCT) sequestration inside cilia, indicating that WDR-60 can enter cilia without dynein-2 but requires its activity in retrograde IFT to exit. Thus, we conclude that the NT of WDR-60 can establish links with other components of the IFT machinery to be incorporated into cilia in the absence of dynein-2. This is in agreement with the weaker but persistent interaction between IFT-B components and the human WDR60[Q631*] truncation lacking the DHC2-binding β-propeller domain (Vuolo et al., 2018).

### WDR-60 is required for efficient IFT recycling and cilia-mediated behavior

Two recent studies in human cells showed that WDR60 loss leads to the misplacement of IFT and signaling particles in cilia without greatly affecting axoneme extension (Hamada et al., 2018; Vuolo et al., 2018). Remarkably, in spite of the strong defect in retrograde IFT, we show that the *wdr-60(null)* mutant is still capable of building nearly full-length cilia in *C. elegans*, similar to what was observed in WDR60 knockout (KO) human cells (Vuolo et al., 2018). We also show that WDR-60-deficient cilia are at least partially functional in chemotaxis and osmotic tolerance assays, which contrasts with the strong behavioral defects exhibited by the *xbx-1(null)* mutant, that assembles severely shortened cilia. The mild chemotaxis and osmotic defects that we observe in *wdr-60* mutants likely mirror the signaling defects underlying *WDR60*-associated SRPS (Cossu et al., 2016; Kakar et al., 2018; McInerney-Leo et al., 2013).

We find that WDR-60 disruption results in accumulation of IFT-74, CHE-11(IFT140), and the kinesin-2 subunit KAP-1 inside cilia. However, we also show that WDR-60 disruption has different effects on the distribution profile of these IFT components along cilia: IFT-74 (IFT-B) levels peak at the tip and in the middle-to-distal segment transition (both regions containing microtubule plus-ends) while CHE-11 (IFT-A) and KAP-1 levels peak closer to the ciliary base and the TZ region, which is similar to the distribution profile of the dynein-2 motor itself. This suggests that some IFT subunits might be less efficiently incorporated into retrograde IFT trains in WDR-60-deficient cilia. In agreement, similar differences in IFT component distribution can be discerned in prior studies using WDR60 KO human cells: IFT88 (IFT-B) accumulates inside their cilia peaking mostly at the tip, while the peak levels of accumulated IFT140 and IFT43 (both IFTA) are found closer to the ciliary base (Hamada et al., 2018; Tsurumi et al., 2019; Vuolo et al., 2018).

Interestingly, our results suggest that the WDR-60 NT retains residual activity in IFT, as the accumulation of IFT-A/B particles inside cilia is less pronounced for the *wdr-60(ΔCT)* mutant than for the *wdr-60(null)* mutant. This is consistent with what was observed in WDR60 KO rescue experiments with an equivalent WDR60 β-propeller truncation construct (WDR60[Q631*]) (Vuolo et al., 2018). Apart from the less pronounced IFT accumulations, the loss of the WDR-60 β-propeller leads to defects in dynein-2 incorporation, retrograde IFT kinetics, and cilia-mediated behavior that are similar to those observed upon the complete loss of WDR-60. Thus, our results show that the WDR-60 β-propeller is critical for dynein-2 function.

### Normal TZ integrity and gating function in the absence of WDR-60

A recent study uncovered that the dynein-2 HC CHE-3 is necessary for the stability and gating functions of the TZ (Jensen et al., 2018). Consistent with this, we show that loss of the dynein-2 LIC XBX-1 also impairs the integrity and gating functions of the TZ barrier. In a recent study, Vuolo and colleagues observed mis-localization of the TZ components TMEM67 and RPGRIP1L (MKS-5 ortholog) in a subset of human RPE1 WDR60 KO cells (Vuolo et al., 2018). In contrast, we do not observe any defects in TZ integrity and gating function in *wdr-60* mutants, as judged by the correct localization of multiple TZ components and the complete RPI-2 exclusion from their sensory cilia. We therefore conclude that the integrity and function of the TZ barrier are maintained in the absence of WDR-60 in *C. elegans*. The difference in TZ susceptibility to WDR60 loss may arise from the variations in TZ structure between different types of cilia (Akella et al., 2019; Jana et al., 2018) or potential differences in the minimal threshold of dynein-2 function required for maintaining TZ integrity in each model system.

### Disruption of WDR-60 reduces dynein-2 loading into cilia and the kinetics of retrograde IFT

Prior studies reported that WDR60 loss resulted in the complete disappearance of dynein-2 LIC from the base and axoneme of cilia (Hamada et al., 2018; Vuolo et al., 2018), but no detectable difference was observed in the recruitment of the dynein-2 HC to the ciliary base (Vuolo et al., 2018). Our analyses of endogenously-labeled dynein-2 HC and LIC in *C. elegans* reveals that both subunits still co-localize inside WDR-60-deficient cilia. Importantly, the efficiency of dynein-2 HC recruitment to cilia is substantially decreased in the absence of WDR-60 or its dynein-2-binding β-propeller domain. Given that dynein-2 HC levels in the soma of ciliated sensory neurons are not greatly altered by the loss of WDR-60, our results argue that WDR-60 directly contributes to dynein-2 recruitment to cilia rather than having a significant role in dynein-2 HC stabilization, as is the case for dynein-2 LIC (Taylor et al., 2015; Toropova et al., 2019).

A recent structural study revealed that binding of the WDR60-WDR34 heterodimer contributes to the asymmetric conformation of the autoinhibited dynein-2 complex, and this was proposed to facilitate dynein-2 incorporation onto anterograde IFT trains (Toropova et al., 2017; Toropova et al., 2019). Our quantifications of dynein-2 HC (GFP::CHE-3) signal on anterograde tracks inside *wdr-60* mutant cilia provide the first direct evidence for this model *in vivo*. This conclusion is supported by the fact that the reduction in dynein-2 loading onto anterograde trains (~3fold) is even greater than the reduction in dynein-2 recruitment to cilia (~2fold) in *wdr-60* mutants.

Intriguingly, the observation that a fraction of dynein-2 motors still get incorporated onto anterograde IFT trains indicates that WDR-60 is not the only link that dynein-2 can establish with anterograde trains. This conclusion is compatible with the previously proposed hypothesis that the main contacts between anterograde IFT trains and dynein-2 motors also involve the heavy chain (Toropova et al., 2019).

Although less dynein-2 reaches the cilium tip to then power retrograde IFT, we find that the frequency of retrograde IFT is only mildly affected in the absence of WDR-60. Taken together with the observation that dynein-2 does not accumulate at the ciliary tip, this result argues that WDR-60 is dispensable for dynein-2 activation and the start of retrograde IFT.

### Underpowered retrograde IFT trains fail to push through the TZ barrier to exit cilia in *wdr-60* mutants

Emerging evidence points to an important interplay between IFT-A and the BBsome in regulating the traffic of G protein-coupled receptors in and out of cilia across the TZ, in part by coupling the receptors to IFT trains (reviewed in (Nachury and Mick, 2019)). As a dense gating structure, the TZ has been shown to slow down the passage of motor-powered IFT trains, supporting the notion that this physical barrier offers substantial resistance to the passage of IFT trains (Jensen et al., 2015; Oswald et al., 2018; Prevo et al., 2015). However, little is known about the mechanisms that enable the IFT machinery to pass through the TZ barrier.

Our live imaging analysis shows that WDR-60-deficient retrograde IFT trains are driven by less dynein-2 motors, at severely reduced velocity, and accumulate at the distal side of the TZ. We propose that the accumulation of these underpowered IFT trains reflects their inability to push through the TZ barrier (**Figure 10**). Additional lines of evidence support a model in which a force production threshold needs to be met in order for retrograde IFT to cross the TZ barrier: a) in experiments using purified “untrapped” dimers of GST-dynein-2 motor domains, maximum microtubule gliding velocity can only be achieved after a certain dynein-2 concentration is reached (Toropova et al., 2017); b) the same study showed that DNA origamis mimicking IFT trains were transported less efficiently (less processive runs) when attached to three “untrapped” dynein-2 motor dimers than when attached to seven dimers (Toropova et al., 2017); c) the cooperative action of multiple dynein-2 motors in retrograde IFT has been shown to be capable of generating considerable forces (at least 25 pN) to move against resisting loads (Roberts, 2018; Shih et al., 2013); d) we show in this study that removing the resistance offered by the TZ rescues the exit of the underpowered retrograde IFT trains driven by less dynein-2 motors in *wdr-60* mutants.

**Figure 10.**
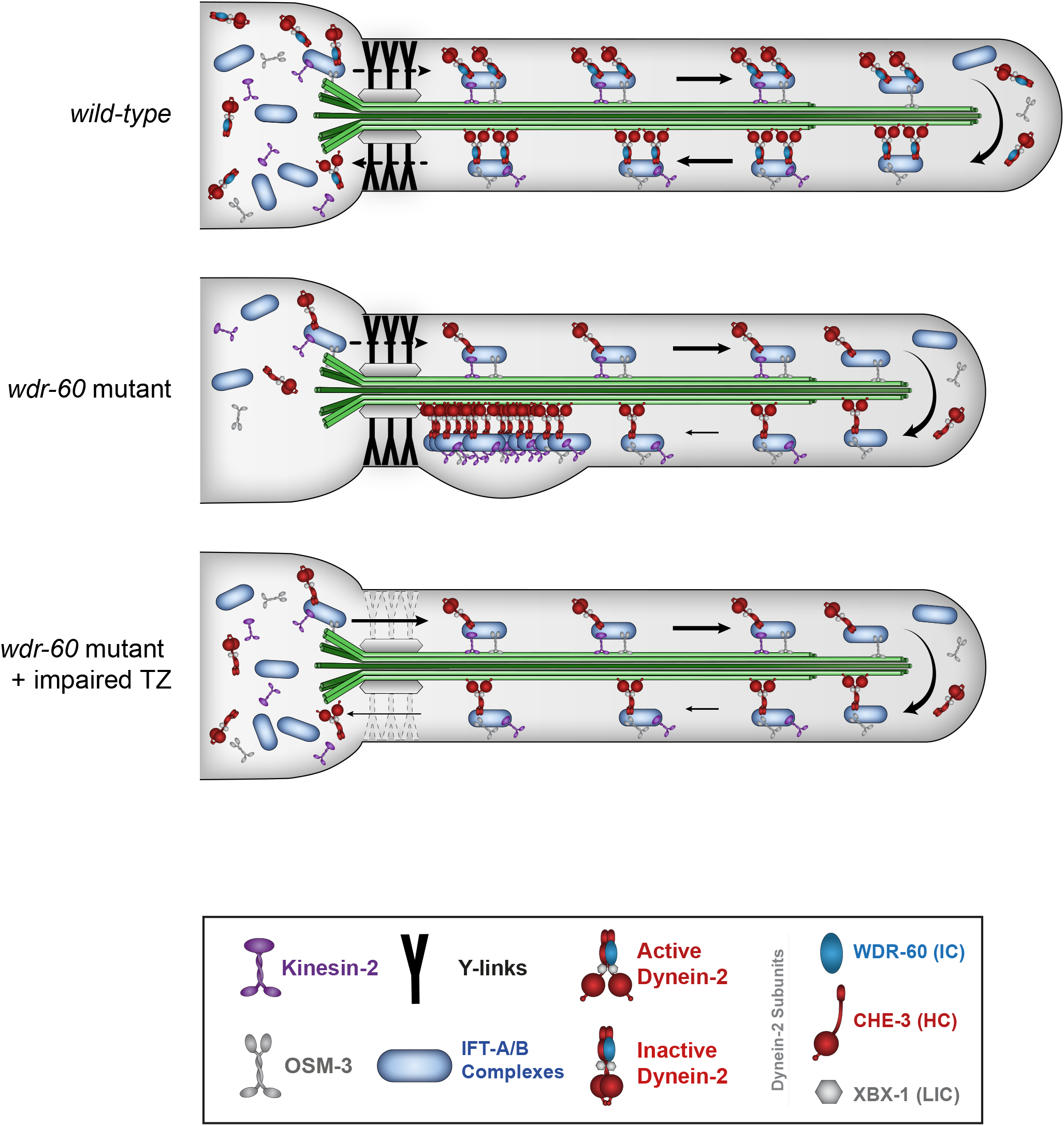
Model for how WDR-60 contributes to efficient dynein-2-mediated retrograde IFT, and the crossing of the TZ to recycle the IFT machinery. **(A)** In wild-type *C. elegans* cilia, kinesin motors carry dynein-2 as a cargo on anterograde IFT trains across the transition zone to enter the cilium compartment and reach the ciliary tip. After rearranged, dynein-2 transports trains in the retrograde direction, crossing the TZ barrier to return them to the base of the cilium, so they can be recycled. **(B)** In the absence of WDR-60, less dynein-2 is recruited and loaded onto anterograde IFT trains to be incorporated into cilia. Consequently, less dynein-2 motors are available at the ciliary tip to power the newly rearranged retrograde trains. These underpowered dynein-2-driven trains move at slower velocities and tend to accumulate at the distal side of the TZ, unable to generate enough force to cross this barrier. **(C)** Disrupting the NPHP module reduces the resistance offered by the TZ to the passage of IFT trains, facilitating the exit of retrograde IFT to the cilium base, compensating for the less efficient WDR-60-deficient dynein-2 trains.

Our findings are also consistent with the “motorized plough” model, which posits that dynein-2 motors remove IFT trains and their cargoes out of the cilium by dragging them while pushing through the TZ barrier (Nachury and Mick, 2019). We note, nonetheless, that we cannot fully exclude the possibility that WDR-60 might also contribute in other ways for retrograde IFT or for ciliary exit.

### The NPHP module offers resistance to dynein-2 passage through the TZ

Our results reveal that disrupting MKS-5, the most upstream TZ assembly factor, can rescue the exit of underpowered retrograde IFT trains from *wdr-60(null)* cilia. We then proceeded to dissect which TZ modules offer resistance to dynein-2 exit by targeting their most upstream components. We find that removal of the NPHP module by disrupting NPHP-4 almost completely rescues the exit of WDR-60-deficient dynein-2 from cilia, while the loss of the MKS module by disrupting MKSR-2 or CEP-290 did not. Given that previous studies showed that the recruitment of NPHP-4 and the assembly of the NPHP module are unaffected by the loss of either CEP-290 or MKSR-2 (Blacque and Sanders, 2014; Li et al., 2016; Schouteden et al., 2015), our findings support a pivotal role for the NPHP module in restricting dynein-2 passage through the TZ. Interestingly, in addition to disrupting the assembly of the NPHP module, NPHP-4 loss has also been shown to reduce the number of Y-links and their densities (Jensen et al., 2015; Lambacher et al., 2016). However, given that the exact contribution of each TZ component to the functionality of Y-links is still not fully understood, it is difficult to directly implicate Y-links in mediating IFT train passage through the TZ.

Consistent with an important role for NPHP-4 and the NPHP module in restricting dynein-2 crossing of the TZ, studies in *C. elegans* and *Chlamydomonas* have shown that NPHP4 loss weakens and permeabilizes the TZ barrier, allowing entry of normally excluded cytoplasmic proteins and reducing the retention of ciliary proteins (Awata et al., 2014; Jauregui et al., 2008; Williams et al., 2011). In agreement with this, we find that removal of NPHP-4 increases the velocity of both anterograde and retrograde IFT trains crossing the TZ, particularly improving the retrograde velocity of underpowered WDR-60-deficient trains exiting cilia. This effect was even comparable with the increased IFT velocity in the TZ region that results from the disruption of MKS-5 in *wdr-60(null)* cilia. This supports the idea that even though NPHP-4 loss does not impair the TZ to the same extent as MKS-5 inhibition, it considerably reduces the resistance offered to IFT trains crossing the TZ barrier.

Interestingly, our results also indicate that even the complete removal of the TZ barrier is not sufficient for clearing out non-motile CHE-3(K2935Q) dynein-2 from cilia. This finding implies that, although reduced, dynein-2 motors powering WDR-60-deficient retrograde IFT trains make an important contribution for the rescue observed upon the disruption of the TZ barrier.

Taken together, our results provide direct evidence that the NPHP module of the TZ offers resistance to the passage of dynein-2-driven IFT trains, and strongly support that dynein-2 motors need to reach a minimal force-generating threshold to power passage of retrograde trains through the TZ barrier to exit cilia.

## Supporting information

Movie S1 - WDR-60 vs GFP-CHE-3

Movie S2 - GFP-CHE-3 in wdr-60 mutants

Movie S3 - CHE-11-mCherry in wdr-60 mutants

Movie S4 - IFT74-GFP in wdr-60 mutants

Movie S5 - GFP-CHE-3 K2539Q in TZ mutants

Movie S6 - CHE-3-GFP in TZ mutants

Movie S7 - CHE-3-GFP in TZ and wdr-60 mutants

## ACKNOWLEDGEMENTS

We thank Dr Alexander Dammermann (Max Perutz) for discussions and Dr José Gama (IBMC/i3S) for critical reading of the manuscript. The authors also thank Drs Alexander Dammermann, Erwin Peterman, Guangshuo Ou, Oliver Blacque, Michel Leroux and Bram Prevo for providing *C. elegans* strains.

This work was financed by FEDER - Fundo Europeu de Desenvolvimento Regional funds through the COMPETE 2020 - Operacional Programme for Competitiveness and Internationalisation (POCI), Portugal 2020, and by Portuguese funds through Fundação para a Ciência e a Tecnologia (FCT) / Ministério da Ciência, Tecnologia e Ensino Superior in the framework of the project POCI-01-0145-FEDER-029471 (PTDC/BIA-BID/29471/2017) to T.J.D.

A.X.C., R.G., C.M.A., and T.J.D. were also supported by the FCT: CEECIND/01967/2017, CEECIND/00333/2017, CEECIND/01985/2018 and CEECIND/00771/2017, respectively. N.V. is a Junior Researcher under the scope of the FCT Transitional Rule DL57/2016.

Some strains were provided by the National Bioresource Project for *C. elegans* and by the *Caenorhabditis* Genetics Center (CGC), which is funded by the NIH Office of Research Infrastructure Programs (P40 OD010440).

## AUTHOR CONTRIBUTIONS

A.R.C. and D.R.R. performed most of the experiments, contributed to the experimental design, analyzed data, helped preparing figures and writing the manuscript. M.J.C. helped with several experiments and figure preparation. N.V. carried out the chemotaxis assay, and C.V. generated some strains. R.G. and A.X.C. provided strains, reagents, equipment and helped with experimental design. T.J.D. and C.M.A. conceived the project, designed and helped with experiments, analyzed and interpreted data, prepared figures, wrote the manuscript and supervised the project. All authors read the manuscript and provided input for the final version.

## SUPPLEMENTAL INFORMATION

**Supplementary file contains Figures S1-S5 and Tables S1-S2.**

**Movie S1 (related to Figure 1).** Live imaging of GFP::CHE-3 (top) and WDR-60::GFP (bottom) undergoing IFT in *C. elegans* phasmid cilia. Images were acquired at 3 fps and playback is set at 15 fps (5x speed). Scale bar, 2 μm. The timer counts mm:ss.

**Movie S2 (related to Figure 3).** Live imaging of GFP::CHE-3 in *C. elegans* phasmid cilia from: control (top); *wdr-60(null)* (center) and *wdr-60(*Δ*CT)* (bottom) strains. Images were acquired at 3 fps and playback is set at 15 fps (5x speed). Scale bar, 2 μm. The timer counts mm:ss.

**Movie S3 (related to Figure 4).** Live imaging of CHE-11::mCherry in *C. elegans* phasmid cilia from: control (top); *wdr-60(null)* (center) and *wdr-60(*Δ*CT)* (bottom) strains. Scale bar, 2 μm. Images were acquired at 3 fps and playback is set at 15 fps (5x speed). Scale bar, 2 μm. The timer counts mm:ss.

**Movie S4 (related to Figure 4).** Live imaging of IFT-74::GFP in *C. elegans* phasmid cilia from: control (top); *wdr-60(null)* (center) and *wdr-60(*Δ*CT)* (bottom) strains. Images were acquired at 3 fps and playback is set at 15 fps (5x speed). Scale bar, 2 μm. The timer counts mm:ss.

**Movie S5 (related to Figure 8).** Live imaging of GFP::CHE-3(K2935Q) in *C. elegans* phasmid cilia from: control (top); *mks-5(tm3100)* (center) and *nphp-4(tm925)* (bottom) strains. Images were acquired at 3 fps and playback is set at 15 fps (5x speed). Scale bar, 2 μm. The timer counts mm:ss.

**Movie S6 (related to Figure 9).** Live imaging of GFP::CHE-3 in *C. elegans* phasmid cilia from: control (top); *mks-5(tm3100)* (center) and *nphp-4(tm925)* (bottom) strains. Images were acquired at 3 fps and playback is set at 15 fps (5x speed). Scale bar, 2 μm. The timer counts mm:ss.

**Movie S7 (related to Figure 9).** Live imaging of GFP::CHE-3 in *C. elegans* phasmid cilia from: *wdr-60(null)* (top); *wdr-60(null);mks-5(tm3100)* (center) and *wdr-60(null);nphp-4(tm925)* (bottom) strains. Images were acquired at 3 fps and playback is set at 15 fps (5x speed). Scale bar, 2 μm. The timer counts mm:ss.

## MATERIALS AND METHODS

### *Caenorhabditis elegans* maintenance and strain generation

*C. elegans* strains were maintained at 20°C on standard nematode growth medium (NGM) plates seeded with *Escherichia coli* OP50 bacteria, and crossed using standard procedures (Brenner, 1974). Hermaphrodite worms were used in all assays. Mutant genotyping was performed by standard PCR. New loci were engineered by CRISPR-Cas9 using germline microinjection of Cas9 and specific gRNA-expressing constructs, supplemented with homology templates consisting in DNA oligos or large DNA fragments partially single-stranded (Dokshin et al., 2018). The presence of the desired alleles was confirmed by PCR-based genotyping. New strains were outcrossed 4 to 6 times to ensure the absence of potential CRISPR-Cas9 off target mutations. *C. elegans* strains used in this study are listed in **Table S2**.

### Fluorescence imaging

All imaging was carried out using young adult hermaphrodite worms, with the exception of the aging experiments, in which Larval stage 2 and 7/18 days post-adulthood animals were also imaged for comparison. All animals used for imaging were immobilized using 5-10mM Levamisol, and were placed on a 5% agarose pad mounted on a microscope slide.

Cilia imaging to generate signal intensity distribution profiles was carried out using an Axio Observer microscope (Zeiss) equipped with 63x, 1.46 NA objective lens, an Orca Flash 4.0 camera (Hamamatsu), and controlled by ZEN software (Zeiss). Z-stacks were acquired with 0.4 μm between each z-section.

Time-lapse imaging of IFT was performed using an Olympus IX81 (Olympus, UK) inverted microscope coupled to an Andor Revolution XD spinning disk confocal system composed of an iXon^EM^+ DU-897 with 2x port coupler camera (ANDOR Technology, UK), a solid-state laser combiner (ALC-UVP 350i, Andor Technology), and a CSU-X1 confocal scanner (Yokogawa Electric Corporation), controlled by Andor IQ3 software (Andor Technology). 200 frames were recorded for each phasmid cilium at 3 frames/sec (333ms per frame). All imaging was performed in temperature-controlled rooms kept at 20°C.

### Image processing and analyses of live IFT

Z-stack and time-lapse series were processed and analyzed with Fiji software (Image J version 2.0.0-rc-56/1.52 p). Fluorescence signal of ciliary components (such as IFT-74::GFP, CHE-11::mCherry and GFP::CHE-3) were used to measure the length of cilia in wild-type and mutant strains from the center of the base to the ciliary tip. The profile of IFT particle distribution was determined along cilia using fluorescently labeled IFT markers. The signal intensity of each pixel was measured from the base to the tip of each cilium and the relative signal distribution of particles was determined. When indicated, signal intensity values of each profile were normalized to their point of maximum intensity to facilitate comparison between profiles from different mutant combinations, and in these instances the total signal from base to tip was also determined and plotted separately.

Kymographs were generated in ImageJ (NIH) using the KymographClear toolset plugin (version 2.0; http://www.nat.vu.nl/~erwinp/downloads.html; (Mangeol et al., 2016)). To ensure the quality of the data used to analyze IFT dynamics, only cilia without severe malformations and completely visible in single stable focal planes were used to generate kymographs. The KymographDirect software (version 2.1; http://www.nat.vu.nl/~erwinp/downloads.html; (Mangeol et al., 2016)) was used to analyze IFT dynamics, which takes into account background and bleaching automatically, and is able to distinguish and separate anterograde from retrograde IFT. Anterograde tracks of individual IFT particles along the whole cilium length were automatically detected by the program as done in (Mijalkovic et al., 2017), and validated. Given the severity of the retrograde IFT phenotypes in *wdr-60* mutants, the program was unable to robustly detect the retrograde tracks of IFT particles. To account for that, all retrograde tracks (both in mutants and in controls) had to be drawn manually. Continuous IFT tracks drawn in the prior step were then used to automatically determine IFT velocities at different positions along cilia in the KymographDirect. When average velocities were calculated for particular sub-regions of the axoneme, they were sub-grouped as follows: TZ (0 – 1μm), middle segment (1 – 4,5μm) and distal segment (4,5μm – ciliary tip).

To determine the frequency of anterograde and retrograde IFT events (data in Figures 3F, 4I, 9C) we employed the approach used in (Mijalkovic et al., 2017). A vertical straight line was drawn to cross the same position at the ciliary distal segment of each anterograde or retrograde kymograph. The line drawn on each kymograph was then used to generate an intensity profile plot, which allowed to score the number of IFT events in either direction by counting the number of intensity spikes. This quantification reflects the number of distinguishable GFP::CHE-3 particles that move in either direction over time.

The average signal intensities of anterogradely or retrogradely moving GFP::CHE-3 particles were determined using KymographDirect as in (Mijalkovic et al., 2017). The software automatically measured the intensity of all pixels composing each IFT track, including the retrograde tracks that had to be drawn manually, as mentioned above. The pixel values were then averaged to provide a single intensity value, representative of each individual track. Each point of GFP::CHE-3 intensity plotted in Figure 3G corresponds to the average of all of the tracks (a minimum of 15) from a single cilium, for either anterograde or retrograde IFT. At least 15 cilia were used per strain to determine the average GFP::CHE-3 intensity particles moving on either direction.

### Dye Filling

A stock solution containing 8 mg/ml of DiI (1,1’-Dioctadecyl-3,3,3’,3’-Tetramethylindocarbocyanine Perchlorate) in dimethyl formamide was prepared in advance and stored at −20°C. One confluent but not starved plate of worms from the strain to be tested was grown for each experiment and a fresh dilution of DiI solution at 2.5μg/ml was prepared in M9 and kept in the dark covered with aluminum foil. Worms were collected and washed in M9 and incubated in 500μL of working DiI solution for 1 h at room temperature in the dark, with occasional flipping of the tubes. After washed in M9, worms were placed on an NGM seeded plate for 3 h at 20°C to further reduce the background of ingested dye. The neuronal uptake of dye was then examined using the Axio Observer microscope as described above. At least 20 adult hermaphrodite worms were examined for each strain in at least two independent experiments.

### Chemosensing assays

Chemotaxis to distinct attractant compounds was assessed as previously described (Bargmann et al., 1993) using synchronized adults at 20°C. Worms were washed three times in a CTX buffer (1 mM CaCl2, 1 mM MgSO4, 5 mM KH2 PO4 pH 6.0) and about 150-200 worms were placed on the origin point of a 10 cm CTX plate without bacteria. The origin point is equidistant to the chosen attractant (isoamyl alcohol - IA, 10% in absolute ethanol) and to the vehicle (absolute ethanol). To prevent the movement of worms away from the attractant or vehicle region, a solution of 1 M sodium azide was added to these points. Worms were then allowed to freely moved at 20°C for 1 h after which worm distribution was determined and the chemotaxis index calculated as previously described (Bargmann et al., 1993).

### Osmotic Avoidance Assay

Osmotic avoidance assays were performed on NGM non-seeded plates at room temperature (~20°C), following the guidelines of (Sanders et al., 2015). Each repeat was carried out using five young adult hermaphrodite worms of each strain isolated before the experiment. Worms were placed inside of a glycerol ring, with a diameter of approximately 1 cm, freshly prepared with a tube dipped in a 59 % glycerol solution. Worm behavior was immediately monitored for 10 min to determine whether they avoided crossing the glycerol ring or not. Worms that left the ring or stayed in contact with its glycerol border for more than 20 seconds were classified as escapers. Wild-type and *xbx-1 (null)* worms were used as controls.

### Immunoblotting

For immunoblots of *C. elegans* extracts, 4 plates of hermaphrodites reaching confluence were collected, washed three times in M9, and the worm pellet was resuspended in an equal volume of 4xSDS-PAGE sample buffer [250 mM Tris-HCl, pH 6.8, 30% (v/v) glycerol, 8% (w/v) SDS, 200 mM DTT and 0.04% (w/v) bromophenol blue]. The worm suspension in sample buffer was supplemented with ~20 μL of glass beads, incubated for 5 min at 95°C and vortexed for additional 5 min. After boiling and vortexing twice, samples were centrifuged at 20000*xg* for 1 min at room temperature, and supernatants were collected. 20% of each sample was loaded and resolved on a 10% SDS-PAGE gel and transferred to 0.2-μm nitrocellulose membranes (GE Healthcare). Membranes rinsed in PBS/0.1% Tween-20 (PBS-T, pH 7.4), and then blocked with 5% (w/v) nonfat dry milk in PBS-T for 1 h Membranes were incubated with mouse anti-FLAG M2 antibody (1:500, Sigma-Aldrich) or mouse anti-α-tubulin B512 antibody (1:5000, Sigma-Aldrich), overnight at 4°C. On the following day, membranes were washed three times in PBS-T for 10 min each. Membranes were then incubated with secondary antibodies coupled to HRP (Jackson ImmunoResearch, 1:10,000) for 1 h at room temperature and washed three times in PBS-T for 5 min each. Pierce ECL Western Blotting Substrate (Thermo Fisher Scientific) was added to membranes to visualize proteins by chemiluminescence using X-ray film or a Chemidoc station (Bio-Rad). Each immunoblot was repeated several times using samples from independent experiments. The predicted protein sizes were based on the NCBI WDR-60 sequence (NP_001367569.1).

### Data analyses and statistics

Statistical analyses of datasets were performed using GraphPad Prism software. For the majority of the experiments, at least 20 worms from different plates were examined for each strain, in at least three independent experiments. Normality tests were carried out to determine whether sample groups followed Gaussian distributions, which dictated the choice between the use of parametric or non-parametric statistical tests. One-way ANOVA, followed up by comparison of the mean of each experimental group with the mean of the control group, was used to analyze parametric datasets, otherwise, we used the non-parametric Kruskal-Wallis test. For single comparison statistical analyses of parametric datasets, the Student’ *t*-test was used, while the Mann-Whitney test was used for non-parametric data. Differences were considered significant at *P* values below 5% (\**P*≤0.05; \*\**P*≤0.01; \*\*\**P*≤0.001; \*\*\*\**P*≤0.0001). XY velocity and intensity distribution graphs are shown as mean ±SEM. Graphs in columns are shown as mean ±SD.

## REFERENCES

Akella, J.S., M. Silva, N.S. Morsci, K.C. Nguyen, W.J. Rice, D.H. Hall, and M.M. Barr. 2019. Cell type-specific structural plasticity of the ciliary transition zone in C. elegans. Biol Cell. 111:95–107.

Asante, D., N.L. Stevenson, and D.J. Stephens. 2014. Subunit composition of the human cytoplasmic dynein-2 complex. J Cell Sci. 127:4774–4787.

Awata, J., S. Takada, C. Standley, K.F. Lechtreck, K.D. Bellve, G.J. Pazour, K.E. Fogarty, and G.B. Witman. 2014. NPHP4 controls ciliary trafficking of membrane proteins and large soluble proteins at the transition zone. J Cell Sci. 127:4714–4727.

Bae, Y.K., and M.M. Barr. 2008. Sensory roles of neuronal cilia: cilia development, morphogenesis, and function in C. elegans. Front Biosci. 13:5959–5974.

Bargmann, C.I., E. Hartwieg, and H.R. Horvitz. 1993. Odorant-selective genes and neurons mediate olfaction in C. elegans. Cell. 74:515–527.

Blacque, O.E., E.A. Perens, K.A. Boroevich, P.N. Inglis, C. Li, A. Warner, J. Khattra, R.A. Holt, G. Ou, A.K. Mah, S.J. McKay, P. Huang, P. Swoboda, S.J. Jones, M.A. Marra, D.L. Baillie, D.G. Moerman, S. Shaham, and M.R. Leroux. 2005. Functional genomics of the cilium, a sensory organelle. Curr Biol. 15:935–941.

Blacque, O.E., and A.A. Sanders. 2014. Compartments within a compartment: what C. elegans can tell us about ciliary subdomain composition, biogenesis, function, and disease. Organogenesis. 10:126–137.

Brenner, S. 1974. The genetics of Caenorhabditis elegans. Genetics. 77:71–94.

Cornils, A., A.K. Maurya, L. Tereshko, J. Kennedy, A.G. Brear, V. Prahlad, O.E. Blacque, and P. Sengupta. 2016. Structural and Functional Recovery of Sensory Cilia in C. elegans IFT Mutants upon Aging. PLoS Genet. 12:e1006325.

Cossu, C., F. Incani, M.L. Serra, A. Coiana, G. Crisponi, L. Boccone, and M.C. Rosatelli. 2016. New mutations in DYNC2H1 and WDR60 genes revealed by whole-exome sequencing in two unrelated Sardinian families with Jeune asphyxiating thoracic dystrophy. Clin Chim Acta. 455:172–180.

Dagoneau, N., M. Goulet, D. Genevieve, Y. Sznajer, J. Martinovic, S. Smithson, C. Huber, G. Baujat, E. Flori, L. Tecco, D. Cavalcanti, A.L. Delezoide, V. Serre, M. Le Merrer, A. Munnich, and V. Cormier-Daire. 2009. DYNC2H1 mutations cause asphyxiating thoracic dystrophy and short rib-polydactyly syndrome, type III. American journal of human genetics. 84:706–711.

Dokshin, G.A., K.S. Ghanta, K.M. Piscopo, and C.C. Mello. 2018. Robust Genome Editing with Short Single-Stranded and Long, Partially Single-Stranded DNA Donors in Caenorhabditis elegans. Genetics. 210:781–787.

Drummond, I.A. 2012. Cilia functions in development. Current opinion in cell biology. 24:24–30.

Garcia-Gonzalo, F.R., and J.F. Reiter. 2017. Open Sesame: How Transition Fibers and the Transition Zone Control Ciliary Composition. Cold Spring Harbor perspectives in biology. 9.

Hamada, Y., Y. Tsurumi, S. Nozaki, Y. Katoh, and K. Nakayama. 2018. Interaction of WDR60 intermediate chain with TCTEX1D2 light chain of the dynein-2 complex is crucial for ciliary protein trafficking. Molecular biology of the cell. 29:1628–1639.

Hou, Y., and G.B. Witman. 2015. Dynein and intraflagellar transport. Experimental cell research. 334:26–34.

Huangfu, D., and K.V. Anderson. 2005. Cilia and Hedgehog responsiveness in the mouse. Proceedings of the National Academy of Sciences of the United States of America. 102:11325–11330.

Huber, C., S. Wu, A.S. Kim, S. Sigaudy, A. Sarukhanov, V. Serre, G. Baujat, K.H. Le Quan Sang, D.L. Rimoin, D.H. Cohn, A. Munnich, D. Krakow, and V. Cormier-Daire. 2013. WDR34 mutations that cause short-rib polydactyly syndrome type III/severe asphyxiating thoracic dysplasia reveal a role for the NF-kappaB pathway in cilia. American journal of human genetics. 93:926–931.

Jana, S.C., S. Mendonca, P. Machado, S. Werner, J. Rocha, A. Pereira, H. Maiato, and M. Bettencourt-Dias. 2018. Differential regulation of transition zone and centriole proteins contributes to ciliary base diversity. Nat Cell Biol. 20:928–941.

Jauregui, A.R., K.C. Nguyen, D.H. Hall, and M.M. Barr. 2008. The Caenorhabditis elegans nephrocystins act as global modifiers of cilium structure. The Journal of cell biology. 180:973–988.

Jensen, V.L., N.J. Lambacher, C. Li, S. Mohan, C.L. Williams, P.N. Inglis, B.K. Yoder, O.E. Blacque, and M.R. Leroux. 2018. Role for intraflagellar transport in building a functional transition zone. EMBO Rep. 19.

Jensen, V.L., C. Li, R.V. Bowie, L. Clarke, S. Mohan, O.E. Blacque, and M.R. Leroux. 2015. Formation of the transition zone by Mks5/Rpgrip1L establishes a ciliary zone of exclusion (CIZE) that compartmentalises ciliary signalling proteins and controls PIP2 ciliary abundance. EMBO J. 34:2537–2556.

Kakar, N., D. Horn, E. Decker, N. Sowada, C. Kubisch, J. Ahmad, G. Borck, and C. Bergmann. 2018. Expanding the phenotype associated with biallelic WDR60 mutations: Siblings with retinal degeneration and polydactyly lacking other features of short rib thoracic dystrophies. Am J Med Genet A. 176:438–442.

Kozminski, K.G., P.L. Beech, and J.L. Rosenbaum. 1995. The Chlamydomonas kinesin-like protein FLA10 is involved in motility associated with the flagellar membrane. The Journal of cell biology. 131:1517–1527.

Kozminski, K.G., K.A. Johnson, P. Forscher, and J.L. Rosenbaum. 1993. A motility in the eukaryotic flagellum unrelated to flagellar beating. Proceedings of the National Academy of Sciences of the United States of America. 90:5519–5523.

Lambacher, N.J., A.L. Bruel, T.J. van Dam, K. Szymanska, G.G. Slaats, S. Kuhns, G.J. McManus, J.E. Kennedy, K. Gaff, K.M. Wu, R. van der Lee, L. Burglen, D. Doummar, J.B. Riviere, L. Faivre, T. Attie-Bitach, S. Saunier, A. Curd, M. Peckham, R.H. Giles, C.A. Johnson, M.A. Huynen, C. Thauvin-Robinet, and O.E. Blacque. 2016. TMEM107 recruits ciliopathy proteins to subdomains of the ciliary transition zone and causes Joubert syndrome. Nat Cell Biol. 18:122–131.

Li, C., V.L. Jensen, K. Park, J. Kennedy, F.R. Garcia-Gonzalo, M. Romani, R. De Mori, A.L. Bruel, D. Gaillard, B. Doray, E. Lopez, J.B. Riviere, L. Faivre, C. Thauvin-Robinet, J.F. Reiter, O.E. Blacque, E.M. Valente, and M.R. Leroux. 2016. MKS5 and CEP290 Dependent Assembly Pathway of the Ciliary Transition Zone. PLoS Biol. 14:e1002416.

Mangeol, P., B. Prevo, and E.J. Peterman. 2016. KymographClear and KymographDirect: two tools for the automated quantitative analysis of molecular and cellular dynamics using kymographs. Molecular biology of the cell. 27:1948–1957.

May, S.R., A.M. Ashique, M. Karlen, B. Wang, Y. Shen, K. Zarbalis, J. Reiter, J. Ericson, and A.S. Peterson. 2005. Loss of the retrograde motor for IFT disrupts localization of Smo to cilia and prevents the expression of both activator and repressor functions of Gli. Developmental biology. 287:378–389.

McInerney-Leo, A.M., M. Schmidts, C.R. Cortes, P.J. Leo, B. Gener, A.D. Courtney, B. Gardiner, J.A. Harris, Y. Lu, M. Marshall, U.K. Consortium, P.J. Scambler, P.L. Beales, M.A. Brown, A. Zankl, H.M. Mitchison, E.L. Duncan, and C. Wicking. 2013. Short-rib polydactyly and Jeune syndromes are caused by mutations in WDR60. American journal of human genetics. 93:515–523.

Merrill, A.E., B. Merriman, C. Farrington-Rock, N. Camacho, E.T. Sebald, V.A. Funari, M.J. Schibler, M.H. Firestein, Z.A. Cohn, M.A. Priore, A.K. Thompson, D.L. Rimoin, S.F. Nelson, D.H. Cohn, and D. Krakow. 2009. Ciliary Abnormalities Due to Defects in the Retrograde Transport Protein DYNC2H1 in Short-Rib Polydactyly Syndrome. American journal of human genetics. 84:542–549.

Mijalkovic, J., B. Prevo, F. Oswald, P. Mangeol, and E.J. Peterman. 2017. Ensemble and singlemolecule dynamics of IFT dynein in Caenorhabditis elegans cilia. Nature communications. 8:14591.

Mikami, A., S.H. Tynan, T. Hama, K. Luby-Phelps, T. Saito, J.E. Crandall, J.C. Besharse, and R.B. Vallee. 2002. Molecular structure of cytoplasmic dynein 2 and its distribution in neuronal and ciliated cells. J Cell Sci. 115:4801–4808.

Nachury, M.V., and D.U. Mick. 2019. Establishing and regulating the composition of cilia for signal transduction. Nat Rev Mol Cell Biol. 20:389–405.

Niceta, M., K. Margiotti, M.C. Digilio, V. Guida, A. Bruselles, S. Pizzi, A. Ferraris, L. Memo, N. Laforgia, M.L. Dentici, F. Consoli, I. Torrente, V.L. Ruiz-Perez, B. Dallapiccola, B. Marino, A. De Luca, and M. Tartaglia. 2018. Biallelic mutations in DYNC2LI1 are a rare cause of Ellis-van Creveld syndrome. Clinical genetics. 93:632–639.

Oswald, F., B. Prevo, S. Acar, and E.J.G. Peterman. 2018. Interplay between Ciliary Ultrastructure and IFT-Train Dynamics Revealed by Single-Molecule Super-resolution Imaging. Cell Rep. 25:224–235.

Patel-King, R.S., R.M. Gilberti, E.F. Hom, and S.M. King. 2013. WD60/FAP163 is a dynein intermediate chain required for retrograde intraflagellar transport in cilia. Molecular biology of the cell. 24:2668–2677.

Pazour, G.J., B.L. Dickert, and G.B. Witman. 1999. The DHC1b (DHC2) isoform of cytoplasmic dynein is required for flagellar assembly. J Cell Biol. 144:473–481.

Porter, M.E., R. Bower, J.A. Knott, P. Byrd, and W. Dentler. 1999. Cytoplasmic dynein heavy chain 1b is required for flagellar assembly in Chlamydomonas. Mol Biol Cell. 10:693–712.

Prevo, B., P. Mangeol, F. Oswald, J.M. Scholey, and E.J. Peterman. 2015. Functional differentiation of cooperating kinesin-2 motors orchestrates cargo import and transport in C. elegans cilia. Nat Cell Biol. 17:1536–1545.

Prevo, B., J.M. Scholey, and E.J.G. Peterman. 2017. Intraflagellar transport: mechanisms of motor action, cooperation, and cargo delivery. FEBS J. 284:2905–2931.

Rana, A.A., J.P. Barbera, T.A. Rodriguez, D. Lynch, E. Hirst, J.C. Smith, and R.S. Beddington. 2004. Targeted deletion of the novel cytoplasmic dynein mD2LIC disrupts the embryonic organiser, formation of the body axes and specification of ventral cell fates. Development. 131:4999–5007.

Roberts, A.J. 2018. Emerging mechanisms of dynein transport in the cytoplasm versus the cilium. Biochem Soc Trans. 46:967–982.

Rompolas, P., L.B. Pedersen, R.S. Patel-King, and S.M. King. 2007. Chlamydomonas FAP133 is a dynein intermediate chain associated with the retrograde intraflagellar transport motor. J Cell Sci. 120:3653–3665.

Sanders, A.A., J. Kennedy, and O.E. Blacque. 2015. Image analysis of Caenorhabditis elegans ciliary transition zone structure, ultrastructure, molecular composition, and function. Methods Cell Biol. 127:323–347.

Schafer, J.C., C.J. Haycraft, J.H. Thomas, B.K. Yoder, and P. Swoboda. 2003. XBX-1 encodes a dynein light intermediate chain required for retrograde intraflagellar transport and cilia assembly in Caenorhabditis elegans. Molecular biology of the cell. 14:2057–2070.

Scheidel, N., and O.E. Blacque. 2018. Intraflagellar Transport Complex A Genes Differentially Regulate Cilium Formation and Transition Zone Gating. Curr Biol. 28:3279–3287 e3272.

Schmidts, M., Y. Hou, C.R. Cortes, D.A. Mans, C. Huber, K. Boldt, M. Patel, J. van Reeuwijk, J.M. Plaza, S.E. van Beersum, Z.M. Yap, S.J. Letteboer, S.P. Taylor, W. Herridge, C.A. Johnson, P.J. Scambler, M. Ueffing, H. Kayserili, D. Krakow, S.M. King, Uk10K, P.L. Beales, L. Al-Gazali, C. Wicking, V. Cormier-Daire, R. Roepman, H.M. Mitchison, and G.B. Witman. 2015. TCTEX1D2 mutations underlie Jeune asphyxiating thoracic dystrophy with impaired retrograde intraflagellar transport. Nature communications. 6:7074.

Schmidts, M., J. Vodopiutz, S. Christou-Savina, C.R. Cortes, A.M. McInerney-Leo, R.D. Emes, H. H. Arts, B. Tuysuz, J. D’Silva, P.J. Leo, T.C. Giles, M.M. Oud, J.A. Harris, M. Koopmans, M. Marshall, N. Elcioglu, A. Kuechler, D. Bockenhauer, A.T. Moore, L.C. Wilson, A.R. Janecke, M.E. Hurles, W. Emmet, B. Gardiner, B. Streubel, B. Dopita, A. Zankl, H. Kayserili, P.J. Scambler, M.A. Brown, P.L. Beales, C. Wicking, Uk10K, E.L. Duncan, and H.M. Mitchison. 2013. Mutations in the gene encoding IFT dynein complex component WDR34 cause Jeune asphyxiating thoracic dystrophy. American journal of human genetics. 93:932–944.

Schouteden, C., D. Serwas, M. Palfy, and A. Dammermann. 2015. The ciliary transition zone functions in cell adhesion but is dispensable for axoneme assembly in C. elegans. The Journal of cell biology. 210:35–44.

Shih, S.M., B.D. Engel, F. Kocabas, T. Bilyard, A. Gennerich, W.F. Marshall, and A. Yildiz. 2013. Intraflagellar transport drives flagellar surface motility. Elife. 2:e00744.

Swoboda, P., H.T. Adler, and J.H. Thomas. 2000. The RFX-type transcription factor DAF-19 regulates sensory neuron cilium formation in C. elegans. Mol Cell. 5:411–421.

Taylor, S.P., T.J. Dantas, I. Duran, S. Wu, R.S. Lachman, C. University of Washington Center for Mendelian Genomics, S.F. Nelson, D.H. Cohn, R.B. Vallee, and D. Krakow. 2015. Mutations in DYNC2LI1 disrupt cilia function and cause short rib polydactyly syndrome. Nature communications. 6:7092.

Toropova, K., M. Mladenov, and A.J. Roberts. 2017. Intraflagellar transport dynein is autoinhibited by trapping of its mechanical and track-binding elements. Nature structural & molecular biology. 24:461–468.

Toropova, K., R. Zalyte, A.G. Mukhopadhyay, M. Mladenov, A.P. Carter, and A.J. Roberts. 2019. Structure of the dynein-2 complex and its assembly with intraflagellar transport trains. Nature structural & molecular biology. 26:823–829.

Tsurumi, Y., Y. Hamada, Y. Katoh, and K. Nakayama. 2019. Interactions of the dynein-2 intermediate chain WDR34 with the light chains are required for ciliary retrograde protein trafficking. Molecular biology of the cell. 30:658–670.

Vuolo, L., N.L. Stevenson, K.J. Heesom, and D.J. Stephens. 2018. Dynein-2 intermediate chains play crucial but distinct roles in primary cilia formation and function. Elife. 7.

Vuolo, L., N.L. Stevenson, A.G. Mukhopadhyay, A.J. Roberts, and D.J. Stephens. 2020. Cytoplasmic dynein-2 at a glance. J Cell Sci. 133.

Webb, S., A.G. Mukhopadhyay, and A.J. Roberts. 2020. Intraflagellar transport trains and motors: Insights from structure. Semin Cell Dev Biol. 107:82–90.

Wicks, S.R., C.J. de Vries, H.G. van Luenen, and R.H. Plasterk. 2000. CHE-3, a cytosolic dynein heavy chain, is required for sensory cilia structure and function in Caenorhabditis elegans. Developmental biology. 221:295–307.

Williams, C.L., C. Li, K. Kida, P.N. Inglis, S. Mohan, L. Semenec, N.J. Bialas, R.M. Stupay, N. Chen, O.E. Blacque, B.K. Yoder, and M.R. Leroux. 2011. MKS and NPHP modules cooperate to establish basal body/transition zone membrane associations and ciliary gate function during ciliogenesis. The Journal of cell biology. 192:1023–1041.

Wu, C., J. Li, A. Peterson, K. Tao, and B. Wang. 2017. Loss of dynein-2 intermediate chain Wdr34 results in defects in retrograde ciliary protein trafficking and Hedgehog signaling in the mouse. Hum Mol Genet. 26:2386–2397.

Yi, P., W.J. Li, M.Q. Dong, and G. Ou. 2017. Dynein-Driven Retrograde Intraflagellar Transport Is Triphasic in C. elegans Sensory Cilia. Curr Biol. 27:1448–1461 e1447.

