## Supplemental Figures and Tables for "WDR60-mediated dynein-2 loading into cilia powers retrograde IFT and transition zone crossing"

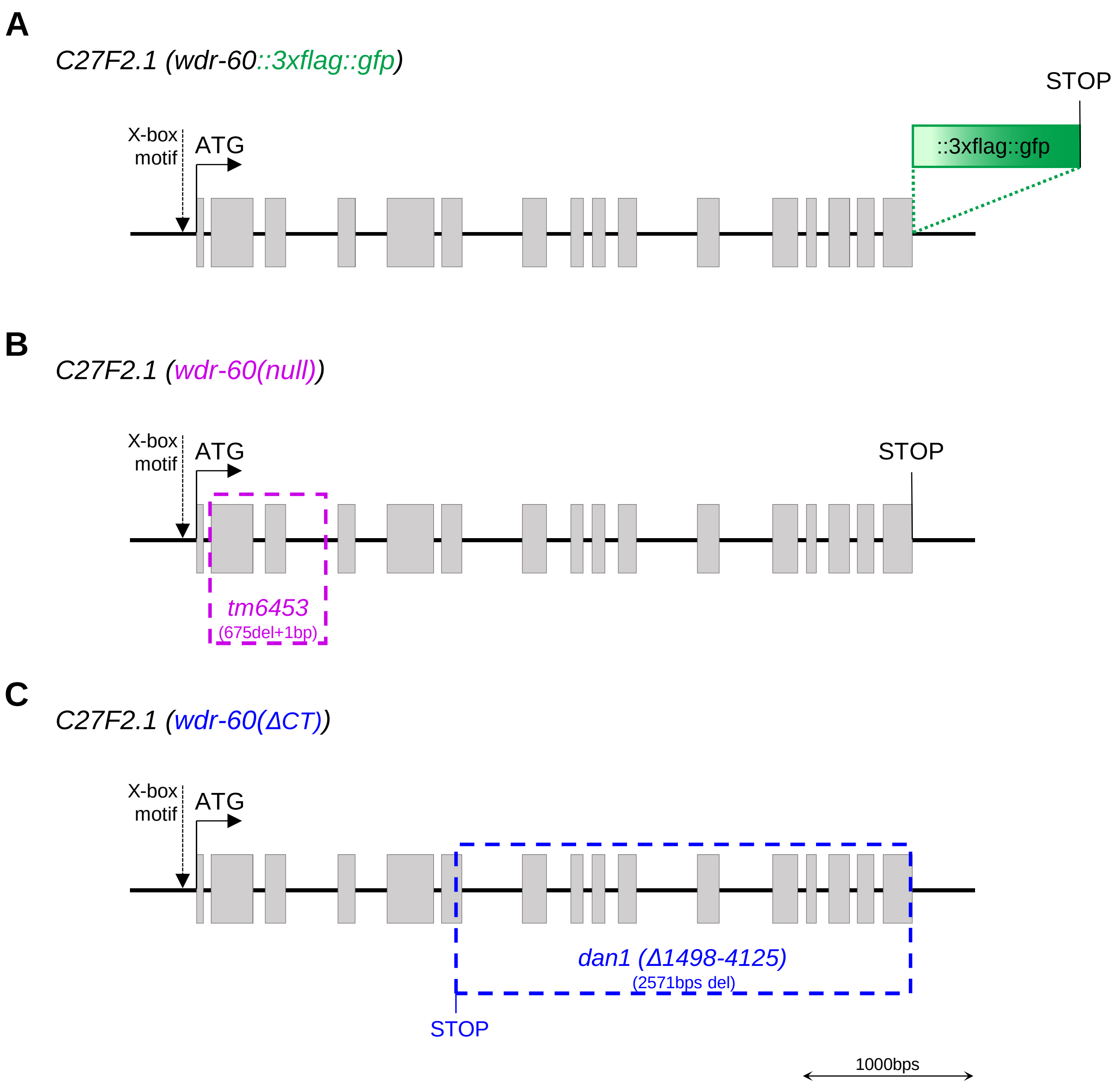


**Figure S1 (related to Figure 1). Schematic of the *C27F2.1* locus organization in the *C. elegans* genome.** Predicted *wdr-60* exons (in grey boxes), including the start and stop codons, and the X-box motif according to (Blacque *et al*., 2005).

**(A)** Knock-in insertion of the *::3xflag::gfp* sequence at the 3'end of the *wdr-60* genomic sequence (in-frame with the WDR-60 coding sequence).

**(B)** Representation of the *wdr-60(tm6453)* null allele.

**(C)** Representation of the *wdr-60(∆CT)* allele (also named as *dan1(∆1498-4125 bps))* corresponding to a WDR-60 truncation of the C-terminal β-propeller domain (∆288-668 aa).

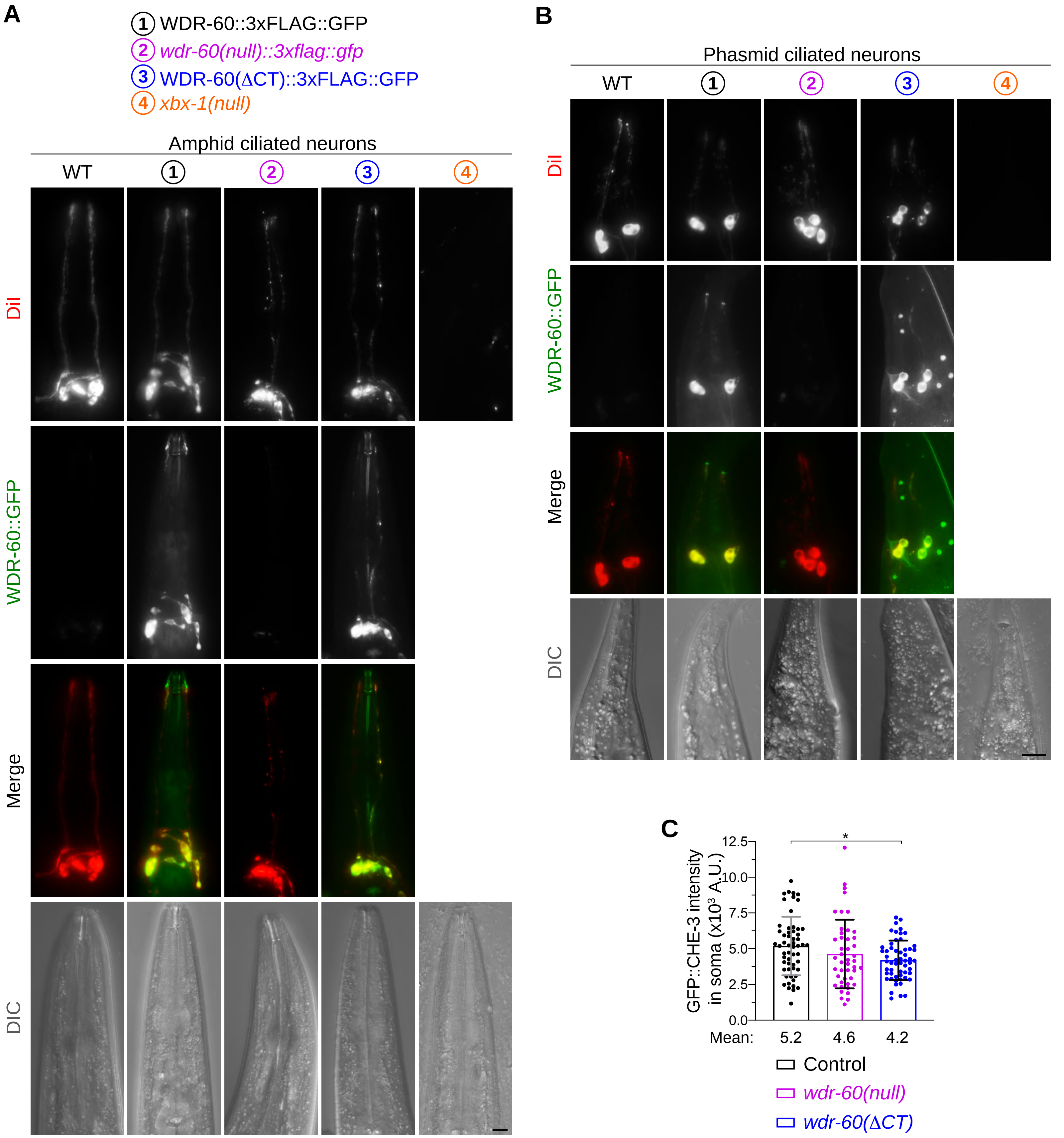


**Figure S2 (related to Figures 2, 3). WDR-60::3xFLAG::GFP expression is restricted to ciliated sensory neurons.**

**(A, B)** Endogenously-tagged WDR-60::3xFLAG::GFP and WDR-60(ΔCT)::3xFLAG::GFP are expressed in the same **(A)** amphid and **(B)** phasmid ciliated neurons that incorporate the DiI lipophilic dye (in red). The *wdr-60(null)::3xflag::gfp* strain has no detectable GFP signal in its neurons; however, these are still able to uptake the DiI dye. Scale bars, 10 μm.

**(C)** GFP::CHE-3 intensity in soma of phasmid sensory neurons of unlabeled *wdr-60* mutants (N≥ 45 somas).

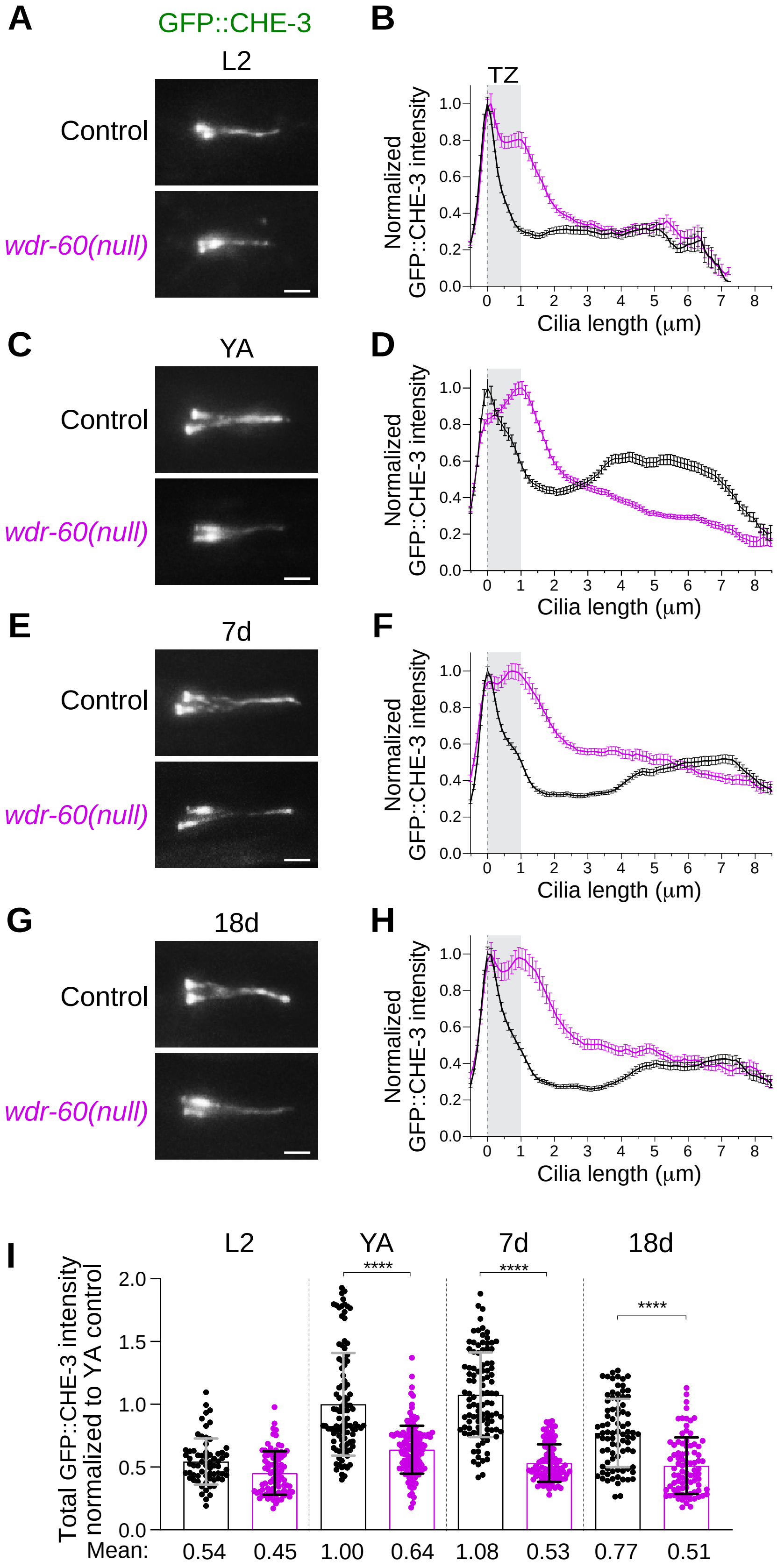


**Figure S3 (related to Figure 3). WDR-60-associated phenotypes become worse as the animals develop to adulthood, and do not improve with aging.**

**(A,C,E,G)** GFP::CHE-3-expressing phasmid cilia of wild-type and *wdr-60(null)* mutants at several stages of development and aging. Scale bar, 2 μm.

**(B,D,F,H)** Corresponding distribution of GFP::CHE-3 signal intensity along cilia. Gray rectangles highlight the TZ, as previously defined.

**(I)** Column graphs showing GFP::CHE-3 total intensity inside cilia of wild-type and *wdr-60(null)* mutants (N≥ 66 cilia). L2: L2 stage, YA: young adult, 7d: 7 days after adulthood, 18d: 18 days after adulthood.

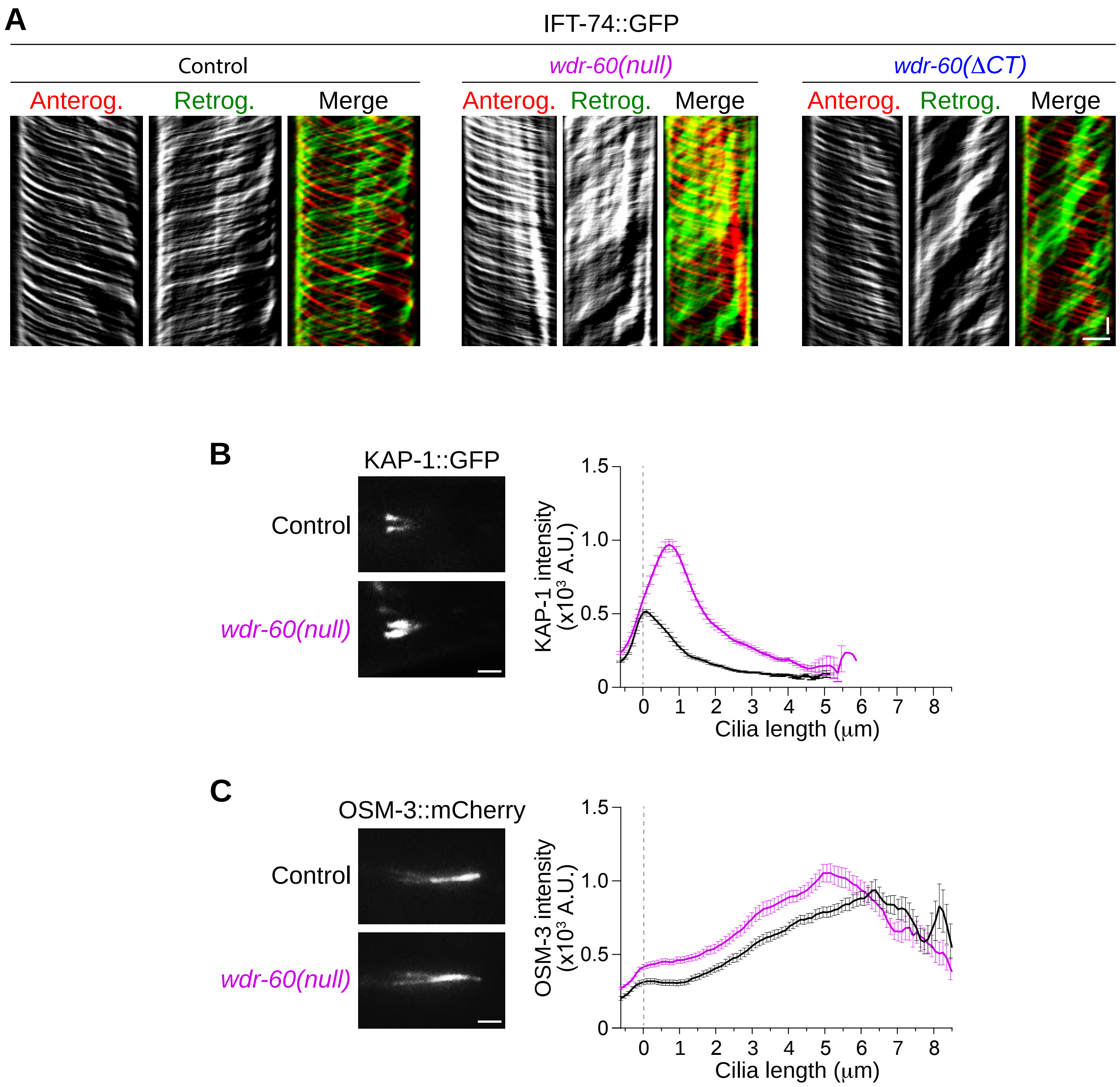


**Figure S4 (related to Figure 4). Loss of WDR-60 results in a severe accumulation of KAP-1 near the base of cilia but only has a modest effect on the distribution of OSM-3.**

**(A)** IFT-74::GFP kymographs of a phasmid cilia from control and *wdr-60(null)* worms. Single channels of particles moving anterogradely and retrogradely are shown, together with their respective merge.

**(B)** Examples of phasmid cilia from control and *wdr-60(null)* worms expressing KAP-1::GFP and quantification of the signal intensity along cilia (N≥ 60 cilia for each strain). The intensity of KAP-1::GFP in cilia is significantly increased in the *wdr-60(null)* mutant with particles accumulating near the TZ.

**(C)** Examples of phasmid cilia from control and *wdr-60(null)* worms expressing OSM-3::mCherry and quantification of the signal intensity along cilia (N≥ 88 cilia for each strain). OSM-3::mCherry distribution and intensity along the cilium is only slightly altered in the *wdr-60(null)* mutant. Scale bars: (A) vertical 5 sec, horizontal 2 µm, (B,C) 2 µm.

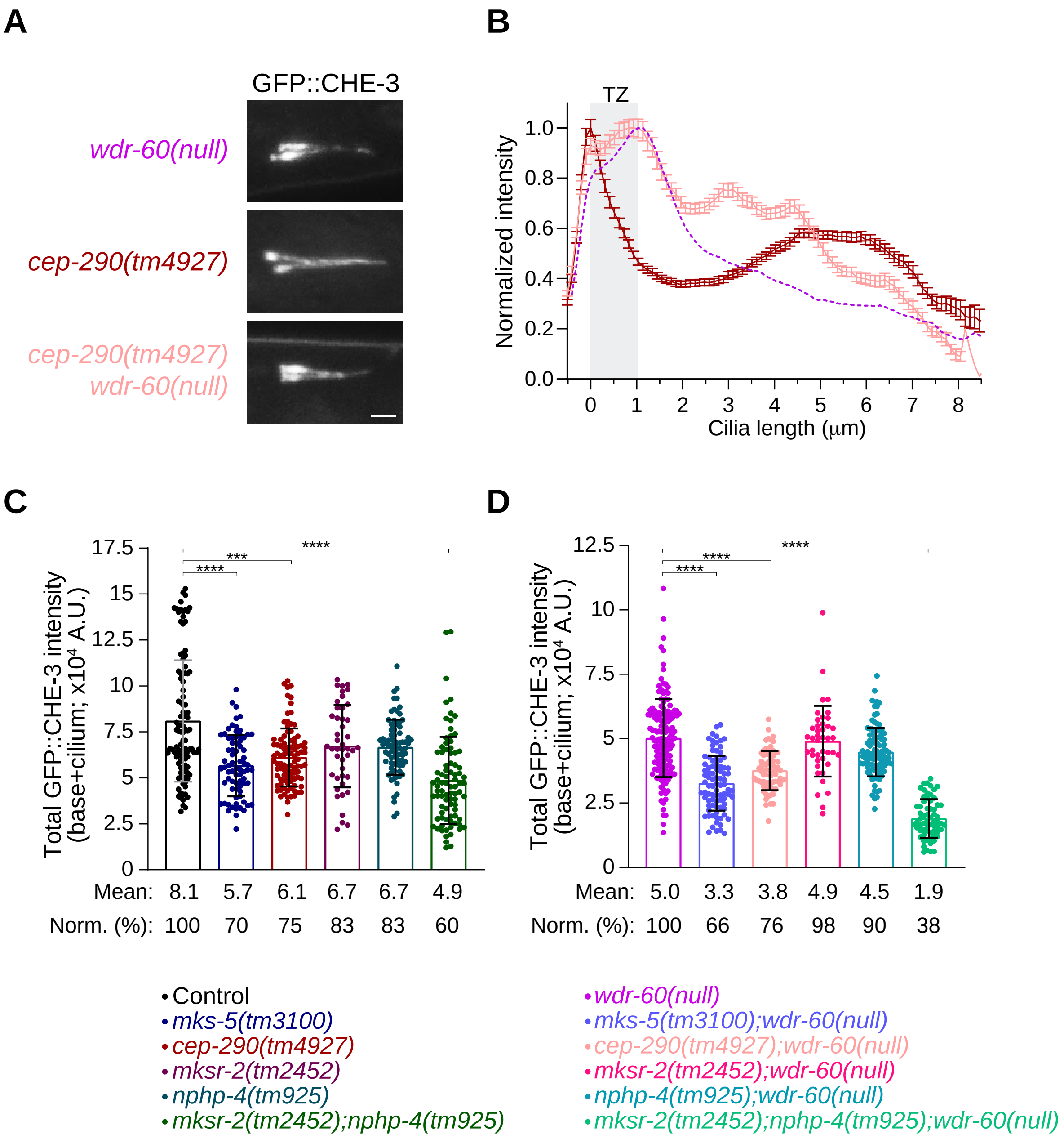


**Figure S5 (related to Figure 7). Disruption of some TZ components changes the average levels of ciliary dynein-2.**

**(A)** Phasmid cilia of the indicated *wdr-60* and *cep-290* mutant genotypes, expressing GFP::CHE-3. Scale bar, 2 μm.

**(B)** Distribution of GFP::CHE-3 signal intensity along cilia from the strains in (A) (N≥ 74 cilia). The gray rectangle highlights the TZ, as previously defined.

**(C,D)** Total GFP::CHE-3 intensity from the base to the tip of cilia from TZ mutants of the indicated genotypes, in **(C)** wild-type *wdr-60* and **(D)** *wdr-60(null)* mutant backgrounds. The cilia quantified here correspond to the same that were used to quantify the GFP::CHE-3 distribution profiles depicted in Figure 7.

**Table S1.** Nomenclature of *C. elegans* proteins that are mentioned in the text, and their corresponding homologs or orthologs in Human.

| | ***C. elegans* proteins** | **Human proteins** | **Other Alias** |
|---|---|---|---|
| **Dynein-2 complex** | CHE-3 | DYNC2H1 | DHC2 |
| | WDR-60 | DYNC2I1 | WDR60 |
| | Unclear | DYNC2I2 | WDR34 |
| | XBX-1 | DYNC2LI1 | LIC3 |
| **IFT Kinesin subunits** | OSM-3 | KIF17 | |
| | KAP-1 | KIFAP3 | |
| **IFT-A Complex** | IFT-43 | IFT43 | |
| | IFT-139 | IFT139/TTC21B | |
| | CHE-11 | IFT140 | |
| **IFT-B Complex** | IFT-74 | IFT74 | |
| | OSM-5 | IFT88 | |
| **Transition Zone components** | MKS-5 | RPGRIP1L/RPGRIP1 | MKS5/NPHP8 |
| | MKSR-1 | B9D1 | |
| | MKSR-2 | B9D2 | |
| | MKS-3 | TMEM67 | MKS3 |
| | MKS-6 | CC2D2A | |
| | TMEM-107 | TMEM107 | JBTS29/MKS13 |
| | CCEP-290/CEP-290 | CEP290 | MKS4/NPHP6/BBS14 |
| | NPHP-4 | NPHP4 | |
| **Transcription factor** | DAF-19 | RFX1/2/3 | |
| **PCMC component** | RPI-2 | RP2 | |
| **Remnant of the degenerated basal body** | HYLS-1 | HYLS1 | |

**Table S2.** List of strains used in this study.

| Strains | Genotype | Short annotation | Reference |
|---|---|---|---|
| N2 | | wild-type | |
| GOU2162 | *che-3 (cas443[gfp::che-3]) I; xbx-1 (cas502[xbx-1::tagRFP::3xFlag]) V* | GFP::CHE-3 + XBX-1::RFP::3xFLAG | Yi et al., 2017 |
| GOU2362 | *ift-74 (cas499[ift-74::gfp]) II* | IFT-74::GFP | Yi et al., 2017 |
| GOU2366 | *che-3(cas511[gfp::che-3(K2935Q)]) I* | GFP::CHE-3(K2935Q) | Yi et al., 2017 |
| JT11069 | *xbx-1 (ok279) V* | *xbx-1 null* | Schafer et al., 2003 |
| DAM459 | *ttTi4391-vieSi23[pAD402; Pnphp-4::gfp::nphp-4cDNA; cb unc-119(+)] I; cxTi10882-vieSi16[pAD390; Phyls-1::mcherry::hyls-1; cb unc-119(+)] IV* | GFP::NPHP-4 + mCherry::HYLS-1 | Schouteden et al., 2015 |
| DAM543 | *cep-290(tm4927)I; mksr-2(tm2452)IV; nphp-4(tm925)V* | Triple mutant of TZ | Schouteden et al., 2015 |
| DAM954 | *vuaSi21 [pBP39; Pmks-6::mks-6::mCherry; cb-unc-119(+)] II* | TZ marker with a mCherry tag | Schouteden et al., 2015 |
| OEB730 | *oqEx500 [tmem-107::GFP + unc-122p::DsRed]* | TMEM-107::GFP | Lambacher et al., 2016 |
| EJP13 | *kap-1(ok676) III; vuaSi1 [pBP20; Pkap-1::kap-1::eGFP; cb-unc-119(+)] IV* | *Pkap-1::kap-1::eGFP* | Prevo et al., 2015 |
| EJP16 | *vuaSi2 [pBP22; Posm-3::osm-3::mCherry; cb-unc-119(+)] II; unc-119(ed3) III; osm-3(p802) IV* | OSM-3::mCherry | Prevo et al., 2015 |
| EJP81 | *vuaSi24 [pBP43; Pche-11::che-11::mCherry; cb-unc-119(+)] II; unc-119(ed3) III; che-11(tm3433) V* | CHE-11::mCherry | Prevo et al., 2015 |
| MX1932 | *nxEx250[rpi-2::gfp; mksr-1::tdtomato; rol-6(su1006)]* | RPI-2::GFP + MKSR-1::tdTomato | Li et al., 2016 |
| AND10 | *che-3 (cas443[gfp::che-3]) I* | GFP::CHE-3 | *this study* |
| AND22 | *ift-74 (cas499[ift-74::gfp]) II; xbx-1 (ok279) V* | IFT-47::GFP + *xbx-1 null* | *this study* |
| AND24 | *che-3 (cas443[gfp::che-3]) I; xbx-1 (ok279) V* | GFP::CHE-3 + *xbx-1 null* | *this study* |
| AND37 | *wdr-60 (tm6453) III* | *wdr-60 null* | *this study* |
| AND38 | *wdr-60 (tm6453) III; ift-74 (cas499[ift-74::gfp]) II* | IFT-47::GFP + *wdr-60 null* | *this study* |
| AND40 | *wdr-60 (tm6453) III; che-3 (cas443[gfp::che-3]) I; xbx-1 (cas502[xbx-1::tagRFP::3xFlag]) V* | GFP::CHE-3 + XBX-1::RFP::3xFLAG + *wdr-60 null* | *this study* |
| AND45 | *wdr-60 (dan1[wdr-60Δ1498_4125]) III* | *wdr-60( ΔCT)* | *this study* |
| AND46 | *wdr-60 (dan1[wdr-60Δ1498_4125]) III; ift-74 (cas499[ift-74::gfp]) II* | IFT-47::GFP + *wdr-60(ΔCT)* | *this study* |
| AND47 | *wdr-60 (dan3[wdr-60:::linker::3xFlag::gfp]) III* | *wdr-60::linker::3xFlag::gfp* | *this study* |
| AND48 | *wdr-60 (dan1[wdr-60Δ1498_4125]) III; che-3 (cas443[gfp::che-3]) I* | GFP::CHE-3 + *wdr-60(ΔCT)* | *this study* |
| AND49 | *wdr-60 (dan1[wdr-60Δ1498_4125]) III; che-3 (cas443[gfp::che-3]) I; xbx-1 (cas502[xbx-1::tagRFP::3xFlag]) V* | GFP::CHE-3 + XBX-1::RFP::3xFLAG + *wdr-60(ΔCT)* | *this study* |
| AND50 | *wdr-60 (dan3[wdr-60:::linker::3xFlag::gfp]) III; xbx-1 (cas502[xbx-1::tagRFP::3xFlag]) V* | wdr-60::linker::3xFlag::gfp + XBX-1::RFP | *this study* |
| AND59 | *wdr-60 (dan3[wdr-60:::linker::3xFlag::gfp]) III; xbx-1 (ok279) V* | *wdr-60::linker::3xFlag::gfp + xbx-1 null* | *this study* |
| AND60 | *wdr-60 (dan1[wdr-60Δ1498_4125]) III; oqEx500 [tmem-107::gfp + unc-122p::DsRed]* | TMEM-107::GFP + *wdr-60(ΔCT)* | *this study* |
| AND66 | *che-3(cas443[gfp::che-3]) I; him-8 (e1489) IV* | GFP::CHE-3 + *him-8 (e1489)* | *this study* |
| AND73 | *wdr-60 (dan4[wdr-60(Δ1498_4125)::linker::3xFlag::gfp]) III* | *wdr-60(ΔCT)::linker::3xFlag::gfp* | *this study* |
| AND79 | *oqEx500 [tmem-107::gfp + unc-122p::DsRed]; xbx-1 (ok279) V* | TMEM-107::GFP + *xbx-1 null* | *this study* |
| AND80 | *wdr-60 (tm6453) III; oqEx500 [tmem-107::gfp + unc-122p::DsRed]; him-8 (e1489) IV* | TMEM-107::GFP + *wdr-60 null* | *this study* |
| AND81 | *wdr-60 (dan5[wdr-60(tm6453)::linker::3xFlag::gfp]) III* | *wdr-60(null)::linker::3xFlag::gfp* | *this study* |
| AND83 | *xbx-1 (ok279) V; wdr-60 (dan4[wdr-60(Δ1498_4125)::linker::3xFlag::gfp]) III* | *wdr-60(ΔCT)::linker::3xFlag::gfp + xbx-1 null* | *this study* |
| AND86 | *che-3 (cas443[gfp::che-3]) I; wdr-60 (tm6453) III* | GFP::CHE-3 + *wdr-60 null* | *this study* |
| AND90 | *wdr-60 (tm6453) III; che-11 (tm3433) V; vuaSi24 [pBP43; Pche-11::che-11::mCherry; cb-unc-119(+)] II* | CHE-11::mCherry + *wdr-60 null* | *this study* |
| AND92 | *wdr-60 (tm6453) III; vuaSi2 [pBP22; Posm-3::osm-3::mCherry; cb-unc-119(+)] II; osm-3(p802) IV* | OSM-3::mCherry + *wdr-60 null* | *this study* |
| AND96 | *che-3 (cas443[gfp::che-3])I; vuaSi21 [pBP39; Pmks-6::mks-6::mCherry; cb-unc-119(+)] II* | GFP::CHE-3 + MKS-6::mCherry | *this study* |
| AND97 | *che-3 (cas443[gfp::che-3])I; mksr-2 (tm2452) IV; nphp-4 (tm925) V* | GFP::CHE-3 + *mksr-2(tm2452) + nphp-4(tm925)* | *this study* |
| AND99 | *wdr-60 (tm6453) III; che-3 (cas443[gfp::che-3]) I; mksr-2 (tm2452) IV; nphp-4 (tm925) V* | GFP::CHE-3 + *wdr-60 null + mksr-2(tm2452) + nphp-4(tm925)* | *this study* |
| AND100 | *wdr-60 (tm6453) III; che-3 (cas443[gfp::che-3]) I; mksr-2 (tm2452) IV* | GFP::CHE-3 + *wdr-60 null + mksr-2(tm2452)* | *this study* |
| AND103 | *xbx-1 (ok279) V; vuaSi21 [pBP39; Pmks-6::mks-6::mCherry; cb-unc-119(+)] II* | MKS-6::mCherry + *xbx-1 null* | *this study* |
| AND104 | *che-3 (cas443[gfp::che-3]) I; wdr-60 (tm6453) III; vuaSi21 [pBP39; Pmks-6::mks-6::mCherry; cb-unc-119(+)] II* | GFP::CHE-3 + MKS-6::mCherry + *wdr-60 null* | *this study* |
| AND110 | *wdr-60 (tm6453) III; vuaSi1 [pBP20; Pkap-1::kap-1::egfp; cb-unc-119(+)] IV* | KAP-1::GFP + *wdr-60 null* | *this study* |
| AND112 | *che-3(cas511[gfp::che-3(K2935Q)]) I; nphp-4 (tm925) V* | GFP::CHE-3(K2935Q) + *nphp-4(tm925)* | *this study* |

| | | | |
|---|---|---|---|
| AND113 | *che-3(cas511[gfp::che-3(K2935Q)]) I; nphp-4 (tm925) V; mksr-2 (tm2452) IV* | GFP::CHE-3(K2935Q) + *nphp-4(tm925)* + *mksr-2(tm2452)* | *this study* |
| AND114 | *che-3(cas511[gfp::che-3(K2935Q)]) I; mksr-2 (tm2452) IV* | GFP::CHE-3(K2935Q) + *mksr-2(tm2452)* | *this study* |
| AND119 | *wdr-60 (dan1[wdr-60Δ1498_4125]) III ; che-11 (tm3433) V; vuaSi24 [pBP43; Pche-11::che-11::mCherry; cb-unc-119(+)] II* | CHE-11::mCherry + *wdr-60(ΔCT)* | *this study* |
| AND122 | *che-3 (cas443[gfp::che-3]) I; wdr-60 (dan1[wdr-60Δ1498_4125]) III ; vuaSi21 [pBP39; Pmks-6::mks-6::mCherry; cb-unc-119(+)] II* | GFP::CHE-3 + MKS-6::mCherry + *wdr-60(ΔCT)* | *this study* |
| AND127 | *vuaSi1 [pBP20; Pkap-1::kap-1::egfp; cb-unc-119(+)] IV* | KAP-1::GFP | *this study* |
| AND129 | *che-3 (cas443[gfp::che-3]) I; mksr-2 (tm2452) IV* | GFP::CHE-3 + *mksr-2(tm2452)* | *this study* |
| AND173 | *che-3(cas443[gfp::che-3])I; nphp-4 (tm925)V* | GFP::CHE-3 + *nphp-4(tm925)* | *this study* |
| AND174 | *che-3(cas443[gfp::che-3]) I; wdr-60 (tm6453) III; nphp-4 (tm925)V* | GFP::CHE-3 + *wdr-60 null* + *nphp-4(tm925)* | *this study* |
| AND184 | *che-3(cas511[gfp::che-3(K2935Q)]) I; mks-5 (tm3100) II* | GFP::CHE-3(K2935Q) + *mks-5(tm3100)* | *this study* |
| AND190 | *che-3(cas443[gfp::che-3]) I; mks-5 (tm3100) II; wdr-60 (tm6453) III* | GFP::CHE-3 + *wdr-60 null* + *mks-5(tm3100)* | *this study* |
| AND194 | *che-3(cas443[gfp::che-3]) I; mks-5 (tm3100) II* | GFP::CHE-3 + *mks-5(tm3100)* | *this study* |
| AND198 | *ttTi4391-vieSi23[pAD402; Pnphp-4::gfp::nphp-4cDNA; cb-unc-119(+)] I; wdr-60 (tm6453) III; cxTi10882-vieSi16[pAD390; Phyls-1::mcherry::hyls-1; cb unc-119(+)] IV* | GFP::NPHP-4 + mCherry::HYLS-1 + *wdr-60 null* | *this study* |
| AND200 | *che-3(cas443[gfp::che-3]) I; wdr-60 (tm6453) III; cxTi10882-vieSi16[pAD390; Phyls-1::mcherry::hyls-1; cb unc-119(+)] IV* | GFP::CHE-3 + mCherry::HYLS-1 + *wdr-60 null* | *this study* |
| AND201 | *che-3(cas443[gfp::che-3]) I; cxTi10882-vieSi16[pAD390; Phyls-1::mcherry::hyls-1; cb unc-119(+)] IV* | GFP::CHE-3 + mCherry::HYLS-1 | *this study* |
| AND203 | *xbx-1(ok279) V; vuaSi24 [pBP43; Pche-11::che-11::mCherry; cb-unc-119(+)] II* | CHE-11::mCherry + *xbx-1 null* | *this study* |
| AND204 | *nxEx250[rpi-2::gfp; mksr-1::tdtomato; rol-6(su1006)]; wdr-60 (dan1[wdr-60Δ1498_4125]) III* | RPI-2::GFP + MKSR-1::tdTomato + *wdr-60(ΔCT)* | *this study* |
| AND207 | *che-3(cas443[gfp::che-3]) I; cep-290 (tm4927) I* | GFP::CHE-3 + *cep-290(tm4927)* | *this study* |
| AND208 | *cep-290 (tm4927) I; che-3(cas443[gfp::che-3]) I; wdr-60 (tm6453) III* | GFP::CHE-3 + *cep-290(tm4927)* + *wdr-60 null* | *this study* |
| AND215 | *ttTi4391-vieSi23[pAD402; Pnphp-4::gfp::nphp-4cDNA; cb-unc-119(+)] I; xbx-1 (ok279) V; cxTi10882-vieSi16[pAD390; Phyls-1::mcherry::hyls-1; cb-unc-119(+)] IV;* | GFP::NPHP-4 + mCherry::HYLS-1 + *xbx-1 null* | *this study* |
| AND216 | *ttTi4391-vieSi23[pAD402; Pnphp-4::gfp::nphp-4cDNA; cb-unc-119(+)] I; cxTi10882-vieSi16[pAD390; Phyls-1::mcherry::hyls-1; cb-unc-119(+)] IV; wdr-60 (dan1[wdr-60Δ1498_4125]) III* | GFP::NPHP-4 + mCherry::HYLS-1 + *wdr-60(ΔCT)* | *this study* |
| AND219 | *nxEx250[rpi-2::gfp; mksr-1::tdtomato; rol-6(su1006)]; wdr-60 (tm6453) III* | RPI-2::GFP + MKSR-1::tdTomato + *wdr-60 null* | *this study* |
| AND222 | *nxEx250[rpi-2::gfp; mksr-1::tdtomato; rol-6(su1006)]; xbx-1(ok279) V* | RPI-2::GFP + MKSR-1::tdTomato + *xbx-1 null* | *this study* |
